# Multigenic resistance to *Xylella fastidiosa* in wild grapes (*Vitis* sps.) and its implications within a changing climate

**DOI:** 10.1101/2022.10.08.511428

**Authors:** Abraham Morales-Cruz, Jonas Aguirre-Liguori, Mélanie Massonnet, Andrea Minio, Mirella Zaccheo, Noe Cochetel, Andrew Walker, Summaira Riaz, Yongfeng Zhou, Dario Cantu, Brandon S. Gaut

**Affiliations:** Dept. of Ecology and Evolutionary Biology, University of California, Irvine, USA; Dept. of Viticulture and Enology; University of California, Davis, USA; Shenzhen Branch, Guangdong Laboratory of Lingnan Modern Agriculture, Genome Analysis Laboratory of the Ministry of Agriculture and Rural Affairs, Agricultural Genomics Institute at Shenzhen, Chinese Academy of Agricultural Sciences, Shenzhen, China; San Joaquin Valley Agricultural Center, United States Dept of Agriculture, Parlier, CA, USA

## Abstract

*Xylella fastidiosa* is a bacterium that infects crops like grapevines, coffee, almonds, citrus and olives, causing economically devastating damage. There is, however, little understanding of the genes that contribute to resistance, the genomic architecture of resistance, and the potential role of climate in shaping resistance, in part because major crops like grapevines (*V. vinifera*) are not resistant to the bacterium. Here we studied a wild grapevine species, *Vitis arizonica*, that segregates for resistance to *X. fastidiosa*. Using genome-wide association, we identified candidate genes that mediate the host response to *X. fastidiosa* infection. We uncovered evidence that resistance requires genes from multiple genomic regions, based on data from breeding populations and from additional *Vitis* species. We also inferred that resistance evolved more than once in the wild, suggesting that wild *Vitis* species may be a rich source for resistance alleles and mechanisms. Finally, resistance in *V. arizonica* was climate dependent, because individuals from low (< 10°C) temperature locations in the wettest quarter were typically susceptible to infection, likely reflecting a lack of pathogen pressure in these climates. Surprisingly, climate was nearly as effective a predictor of resistance phenotypes as some genetic markers. This work underscores that pathogen pressure is likely to increase with climate, but it also provides genetic insight and tools for breeding and transforming resistant crops.

## INTRODUCTION

Climate change is impacting crop yields by shifting temperatures, weather extremes, and water availability (Zhao et al. 2017), thereby affecting the distribution of arable lands (Ramankutty et al. 2002). There is, however, another important effect of climate change, which is the altered distribution of plant pathogens (Velásquez et al. 2018; Burdon and Zhan 2020). One especially prominent pathogen is the bacterium *Xylella fastidiosa. X. fastidiosa* is a generalist that colonizes > 300 plant species (Sicard et al. 2018; European Food Safety Authority (EFSA) 2020), but it is pathogenic on major crops like citrus, coffee, almonds and grapevines (*Vitis vinifera* ssp. *vinifera*). Until recently, *X. fastidiosa* had been limited to the Americas, but human-mediated migration has led to its colonization of Europe, where it causes > ~$100M of damage per year to the olive industry (Schneider et al. 2020). This olive example illustrates that the bacterium is more than a persistent threat in the Americas; it is also an emerging and expanding global threat to Europe, the Middle East (Frem et al. 2021) and beyond (Su et al. 2013). Accordingly, there are urgent needs to better understand the genetic mechanisms of plant resistance (National Research Council et al. 2004), particularly in the wild where both pathogens and hosts evolve (Bartoli and Roux 2017).

Thus far, studies of *X. fastidiosa*-mediated diseases have focused primarily on citrus and on Pierce’s Disease (PD) in domesticated grapevines. In grapevines, PD manifests by colonizing the xylem, leading to vascular blockages and eventual plant death after several years. In the course of infection, PD causes other detrimental symptoms, including marginal leaf necrosis, berry desiccation, irregular maturation of canes and abnormal petiole abscission (Rapicavoli et al. 2018). The bacterium is spread from plant to plant by xylem-feeding insect vectors, which affect the severity and spread disease. The distribution of these insect vectors is being affected by changing climate (Hoddle 2004) and by anthropomorphic activity. One pertinent example is the glassy-winged-sharpshooter (GWSS; *Homalodisca vitripennis*), which was introduced to Southern California in the late 1990s. The GWSS has a higher transmission efficiency compared to native vectors and fueled a large PD outbreak that has permanently altered viticulture in the region.

Although all domesticated grapevines (*V. vinifera* ssp. *vinifera*) are susceptible to PD, some wild relatives of grapevines segregate for PD resistance, likely reflecting the evolution of resistance in regions of persistent *X. fastidiosa* pressure (Ruel and Andrew Walker 2006). Among wild grapevines, *Vitis arizonica* merits particular interest because it exhibits strong resistance to PD and because it contains the only characterized plant locus to segregate for *Xylella fastidiosa* resistance, the *Pierce’s disease resistance 1* (*PdR1*) locus (Krivanek et al. 2006; Riaz et al. 2006). *PdR1* was identified by genetic mapping of a segregating family, defined by simple-sequence-repeat (SSR) markers, and backcrossed into susceptible grapevine cultivars to introduce resistance (Riaz et al. 2009). A recent study utlized BAC sequences of the region to identify two candidate genes for resistance (Agüero et al. 2022). Both genes were canonical leucine-rich receptor (LRR) loci, but neither conferred resistance after single-gene transformation into *V. vinifera* (Agüero et al. 2022). Additional candidates have been identified based on comparative transcriptomics in *V. vinifera* (Zaini et al. 2018), olives (*Olea europaea*)(Giampetruzzi et al. 2016) and citrus (*Citrus reticulata*) (Rodrigues et al. 2013).

Despite the enormous economic impact of *X. fastidiosa* infection, the genomic architecture of resistance has not yet been investigated in any species, and the genomic basis of resistance remains unclear. Here we address this shortcoming by performing genome-wide association (GWA) analyses for *X. fastidiosa* resistance in *V. arizonica*. In addition to identifying several novel candidate genes for resistance in *PdR1* and in other genomic regions, our work begins to fill another surprising gap. Although GWA and similar approaches are commonly used to study disease resistance in crops, surprisingly few studies have focused on the wild relatives of crops (Bartoli and Roux 2017). [One notable exception is the wild relative of soybean, *Glycine soja* (Leamy et al. 2017; Zhang et al. 2017).] This dearth of studies is surprising both because crop wild relatives are a proven and valuable source of resistance genes for crop improvement (Migicovsky and Myles 2017) and because studying resistance in wild samples may provide insights into the evolution of resistance and the ecological and climatic factors that shape resistance (Bartoli and Roux 2017).

In this study, we generate landscape genomic data from a sample of *V. arizonica* from throughout its native range and perform GWA based on a resistance phenotype - i.e., bacterial load after experimental inoculation. In doing so, we identify several genomic regions, including the *PdR1* region, that are associated with resistance, and we identify candidate genes in these regions based on an improved *V. arizonica* reference. We combine GWA with several types of evidence – including population genetic analyses, gene expression assays, comparisons among wild *Vitis* species, investigation of *V. vinifera* cultivars bred for PD resistance and bioclimatic modeling - to address three sets of questions. First, which and how many genic regions contribute to resistance, and what are some of the likely candidate resistance genes within these regions? Second, are these regions implicated in resistance across *Vitis* species and also in cultivars that were specifically bred for PD resistance? What do these inter-species analyses imply about the origin of resistance? Finally, does plant resistance correlate with bioclimate? If so, what might this climatic relationship imply about the potential effects of climate change? Overall, our work provides information about the genetics, evolution and ecology of PD resistance, all of which will help inform strategies to manage an economically damaging and expanding pathogen (Frem et al. 2020).

## RESULTS

### Genome-wide associations for resistance to Pierce’s Disease

We studied the genetics of PD resistance in *V. arizonica* by combining three sources of information: an updated reference genome (accession b40-14, which is homozygous for PD resistance) (Morales-Cruz et al. 2021), whole-genome resequencing data from 167 accessions sampled across the species’ native range (Fig. S1), and previously published phenotypic data about PD resistance on the same set of 167 accessions (Riaz et al. 2020; Morales-Cruz et al. 2021). We used PD resistance as a quantitative variable - i.e., the log-transformed number of colony forming units (CFUs/ml) 12-14 weeks after experimental *X. fastidiosa* inoculations (Table S1). However, following precedence (Riaz et al. 2020), we also characterized individual accessions as resistant if they had *Xylella fastidiosa* concentrations below 13.0 CFUs/ml. Based on this threshold, our sample contained 135 resistant and 32 susceptible individuals, with the susceptible individuals more common in the northern region of the geographic distribution (Fig. 1).

**Fig. 1.**
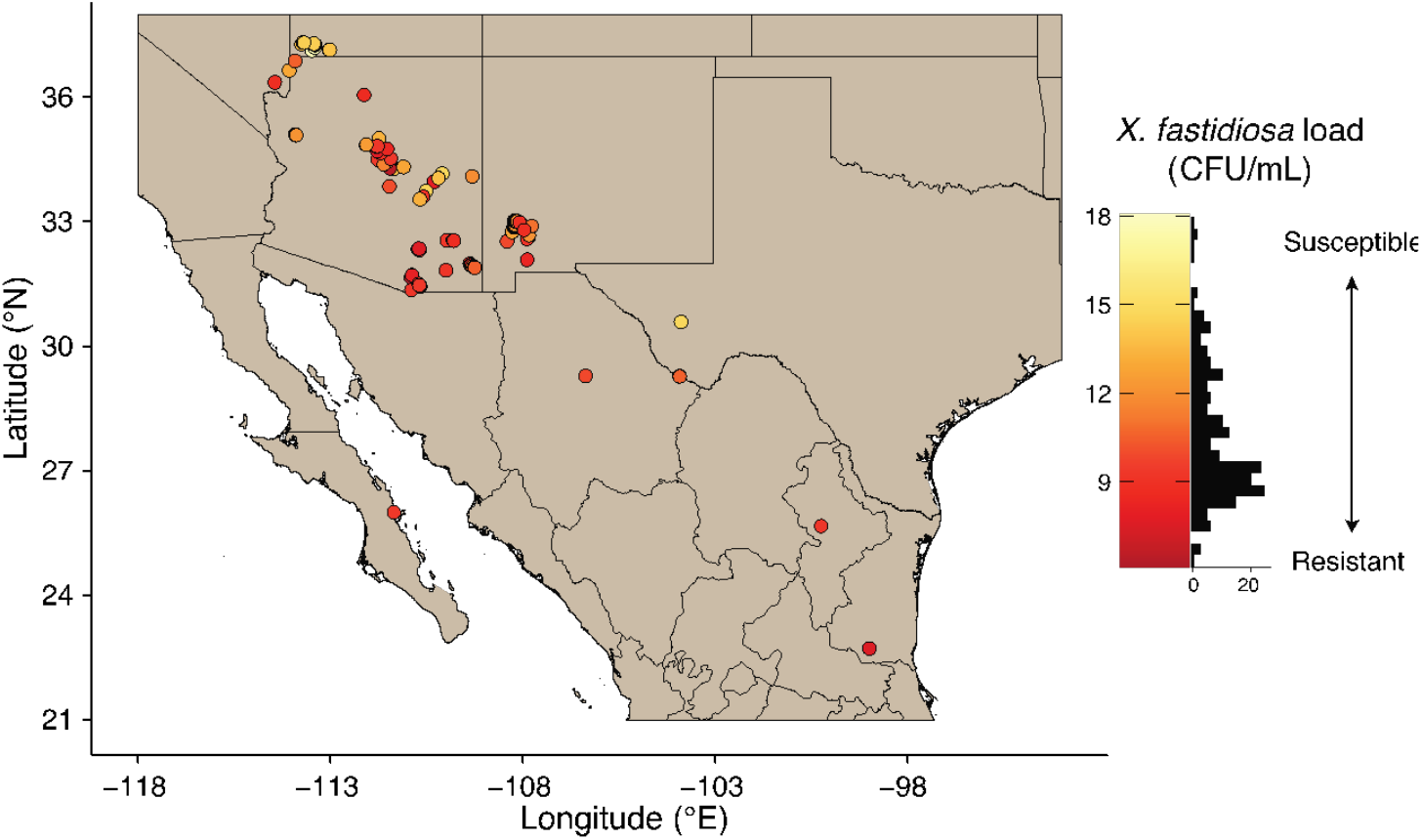
*Vitis arizonica* sampling and phenotypes. A map of the Southwestern United States and Northern Mexico indicates sampling locations of the n=167 *V. arizonica* accessions used in this study. The color of sample locations (circles) are colored according to their resistance phenotype, as measured by bacterial load (CFU/mL). The histogram of phenotypes (in CFU/mL) is to the right of the map.

We first performed genome-wide association (GWA) analyses based on SNP variants. To do so, we mapped resequencing data to the reference haplotype of the phased diploid genome and then tested for associations using two distinct methods that correct for genetic structure (Hao et al. 2021; Caye et al.). On the reference haplotype (hap 1), we identified 74 and 40 associated SNPs (Bonferroni *p* < 0.05) with the two methods, of which 25 were significant with both methods. We used these 25 SNPs to conservatively define eight peaks across five chromosomes (Fig. 2, Figs. S2-S4, Table S2). The most evident peaks were on chromosomes 14 and 15, with the former located between the SSR markers that define the *PdR1* locus. We also called SNPs independently to the second haplotype (hap2) and identified 11 significant SNPs in five peaks (Figs. S5-S7, Table S2). One of these peaks was also on chromosome 14 between the *PdR1* flanking markers.

**Fig. 2.**
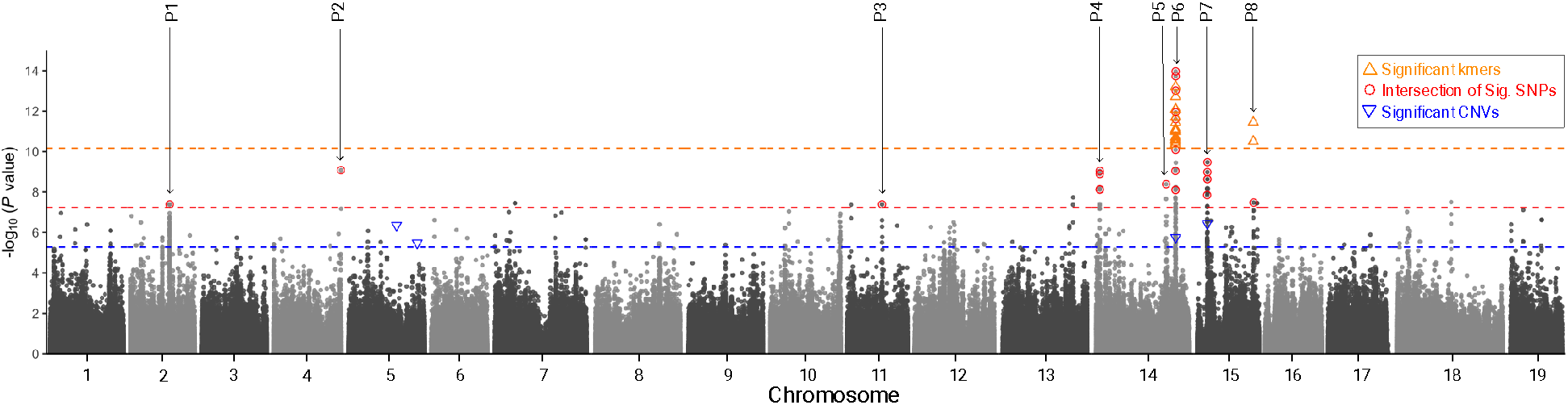
A Manhattan plot of the *V. arizonica* genome showing markers associated with bacterial load. The plot denotes each of the 19 chromosomes for haplotype 1. Each circle represents a SNP with a corresponding p-value, based on EMMAX genome-wide association analysis. The 25 SNPs that were detected in two separate GWA analyzes are circled in red and define the 8 peaks of association, which are numbered as P1, P2, etc., and referred to in the text. In addition to SNPs, the locations of significantly associated kmers and CNVs are provided when they overlap with a SNP-defined peak. The colored horizontal lines represent the cut-off p-values (*P*<0.05, Bonferroni corrected) for the different marker types.

Previous studies have suggested that *PdR1* alleles differ in size among *V. arizonica* accessions (Riaz et al. 2008), suggesting structural variants contribute to resistance. We therefore investigated associations using copy number variants (CNVs), identifying 14,294 CNVs throughout the genome, of which 60 were in the 8 PD-significant peaks (Table S3). The 60 CNVs included 19 deletions and 41 duplications, with means of 1.3 and 3.3 copies. We also performed GWA on the complete CNV set, finding four that were significantly (Bonferroni *p* < 0.05) associated with bacterial load (Fig. 2, Figs. S8-S9). Two of these four CNVs mapped to two SNP-defined peaks: CNV10605 within *PdR1* (mean copy number= 0.88, size= 6kb, R= −0.36, *p*=1.77e-06) and CNV10806 within peak 7 of chromosome 15 (mean copy number = 2.83, size = 17 kb, R= −0.38, p= 3.61e-07). The negative correlations for both CNVs indicated that a higher number of copies had lower bacterial loads and higher PD resistance. Both CNVs had homology to long-terminal repeat transposable elements and so provided few insights into the functional basis of resistance. To further account for potential structural variation among accessions, we also applied GWA to 31bp kmers, using a reference-free approach (Voichek and Weigel 2020) (Fig. 2, Fig. S10). Of 115 significant kmers (Bonferroni *p* < 0.05) (Table S4), 79 mapped to the reference genome (Table S5) and 62 mapped uniquely to either hap1 or hap2. Among the uniquely mapped kmers, 57 of 62 were located on hap1 near *PdR1* and five were on the chromosome 15 peak. Altogether, CNV and kmer analyses corroborated four of the eight SNP-based peaks while confirming *PdR1* as a major locus (Krivanek et al. 2006; Riaz et al. 2006).

We manually reannotated genes under the eight hap1 peaks, using boundaries defined by 100kb windows, since genome-wide LD decayed to background levels (*r*^2^<0.05) within this distance (Fig. S11). The eight peaks included 124 genes, and several had annotations that implied a role in plant immunity (Table S6). For example, peak 4 included a calmodulin-binding gene (g226310) that is involved in the regulation of plant disease response through changes in phytohormone biosynthesis (Levy et al. 2005; Lv et al. 2019), and a “syntaxin of plants 41” gene (g226360) that acts in plant resistance against bacterial pathogens (Kalde et al. 2007). At *PdR1* (peak 5), we identified 7 leucine-rich repeat receptor-like protein (LRR-RLP) genes, one LRR receptor-like protein kinase (RLK) gene, and one lysin motif (LysM) RLK gene, that are commonly involved in pathogen detection and initiate the plant response (Liu et al. 2017). Peak 7 contained two nucleotide-binding site leucine-rich repeat proteins (NBS-LRR; g243780, and g243820) that also detect pathogens and initiate a host response, as well as a Phloem protein 2-like (PP2) protein with antimicrobial properties (Du et al. 2022). We also identified eight genes of interest on chromosome 15 (peak 8). Two of those genes have functional annotations related to phytohormone interactions (g252710, Ethylene-responsive transcription factor CRF4, and g252790, Abscisic Acid Insensitive-like 1 or ABIL 1), and thus may play a role in plant immunity. Another four genes encode acidic endochitinases, which provide defense against fungal pathogens (Samac et al. 1990). Finally, we studied the potential function of the 36 significant kmers that did not map to the reference genome by assembling reads containing the kmers and then aligning assemblies to the NCBI Transcript Reference Sequences (“refseq_rna”). Of the assembled contigs, 80% had high similarity to three specific Receptor-like proteins (RLPs) (“ XM_010648495.2”, “XM_034852027.1” and “XM_019224733.1”). Overall, the set of candidate genes suggest that multiple diverse functions contribute to PD resistance, but with likely involvement of classic disease resistance (R) genes.

We used SSR markers to define the *PdR1* locus as a 361 kb region on hap1 chromosome 14 (with a corresponding 360kb region on hap2), but we further characterized the locus in three ways. First, we evaluated linkage disequilibrium (LD) across chromosome 14. We observed two large blocks (~ 7 Mb in size) in high LD that contained the three PD-significant peaks of chromosome 14 (peaks 4, 5, and 6), even though peaks 5 and 6 were located on opposite ends of the chromosome from peak 4 (Fig. 3A). This striking pattern may simply reflect properties of our sample, but it also suggests that PD-related alleles co-segregate across peaks, implying that additive or epistatic interactions contribute to resistance in nature. Second, we focused on the location of significantly associated *PdR1* markers, which fell into a narrower 103.6kb region containing six genes, three of which were RLPs (Fig. 3B&C).

**Fig. 3.**
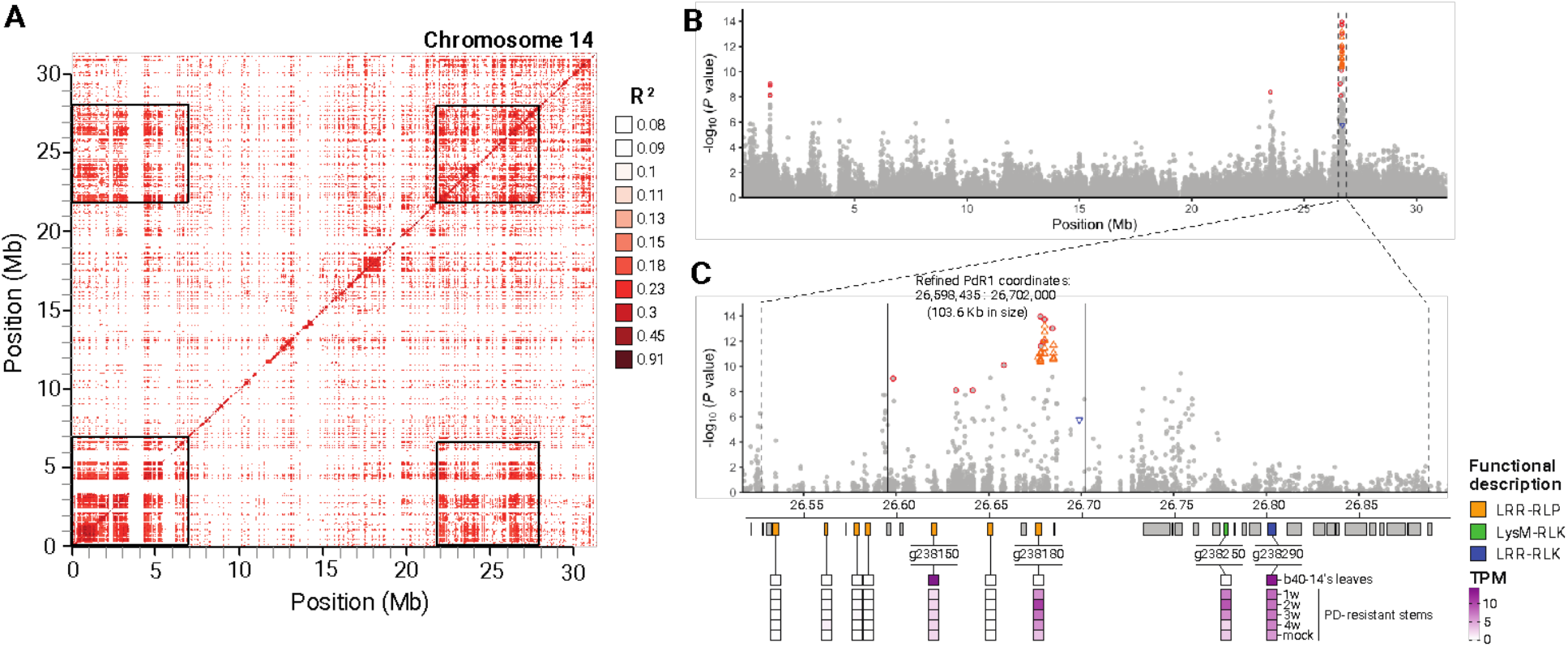
Genetic analyses of the *PdR1* region. **A.** A plot of linkage disequilibrium (LD) across chromosome 14, where darker colors represent higher levels of LD. The two dark squares on the diagonal include GWA peak 4 (on the left, from 0 to 7 Mb on the chromosome) and the GWA peak that corresponds to PdR1 (on the right, from 22 to 28 Mb on the chromosome). The two off-diagonal squares reflect LD between these two distinct regions. **B.** A Manhattan plot of chromosome 14 indicating the locations of peak 4 and the *PdR1* region. **C.**An expanded representation of the *PdR1* region showing the location of significant SNPs (red circles denote SNP significant with two GWA methods), significant kmers (green triangles) and CNVs (blue triangles). The dashed vertical lines represent the 361 kb region defined by SSR markers and the 106kb region defined by the location of significant markers. **D.** A representation of genes in *PdR1* with a summary of gene expression results. Genes are colored if they are related to R genes, with the category of R gene indicated by its color. Expression information shows expression, in transcripts per million (TPM) for leaves, for stems during four stages after infection and for mock controls.

Finally, we assayed gene expression in the region. One RLP (g238150) was expressed in b40-14 leaves, as was a receptor-like-kinase (RLK; g238290) that fell outside the 103.6kb region (Fig. 3C). We also assayed gene expression in three resistant full-sibs that were inoculated with *X. fastidiosa* and a control (water). The stems above the inoculation site were sampled weekly for up to four weeks. Both g238150 and g238290 were expressed at higher levels than the control in at least one weekly stage, although not significantly so (p>0.05). Two additional genes - an RLP (g238180) and an RLK (g239250) - also exhibited this pattern, and g238180 also colocated with several associated kmers (**Fig. 2**). All four of the expressed R genes were also present on the hap2 version of *PdR1*. Importantly, none of these four candidates were the closest homologs of the candidate genes that failed to confer resistance when transformed into *V. vinifera (Agüero et al. 2022*) (see Discussion).

### The genetic basis of resistance in breeding

The complex LD pattern on chromosome 14 suggests that resistance may require genic action from more than one locus - i.e., multigenic (horizontal) resistance. To investigate this possibility, we examined the distribution of kmers across accessions (Table S7). Among the 117 kmers associated with bacterial load, 99 were common among resistant accessions; they were found in 65.0% of resistant plants, on average, but in only 9.5% of susceptible accessions. We labeled these kmers as resistant (R-kmers). In contrast, 16 kmers were detected in 67.2% and 10.1% of susceptible and resistant accessions, on average, suggesting associations with disease susceptibility (S-kmers). Interestingly, 10 of the 16 S-kmers mapped to a region on chromosome 15 that was ~12 kb upstream of a Jasmonic Acid-Amido Synthetase gene (*JAR1*, g252600). Changes in the expression of *JAR1* are associated with a reduction of host defenses (Jiang et al. 2016). We hypothesize that S-kmers are linked to variants that affect the expression of *JAR1* and promote susceptibility to *Xylella fastidiosa*. An important goal for breeding may be to avoid these S-kmers (Zaidi et al. 2018).

We then investigated the genomic content of five resistant cultivars (Ambulo Blanc, Caminante Blanc, Camminare noir, Errante noir and Paseante noir) derived from backcrosses to *V. arizonica* (accession b43-17) (Anon) to test whether the basis of resistance lay solely in *PdR1* or included additional genomic regions. After resequencing the five cultivars, we detected all 99 R-kmers in each cultivar but no S-kmers (Fig. 4B, Table S8). In contrast, a control dataset from four susceptible *V. vinifera* cultivars (Cabernet Sauvignon cl. 08, Chardonnay cl. 04, Zinfandel cl. 03 and Petite Sirah) contained neither R-kmers or S-kmers (Table S8). Although our analyses used a reference (b40-14) that was not the source of PD resistance in backcrossed cultivars (b43-17), we found 56 kmers mapped to b40-14 hap1, 44 to hap2, and 53 to unplaced contigs. Importantly, the hap1 kmers mapped to both *PdR1* (51 kmers) and to peak 8 on chromosome 15 (5 kmers), suggesting these two regions contribute to (and may be necessary for) resistance. As a complementary method, we scanned SNP heterozygosity in resistant cultivars, reasoning that backcrossed regions should be heterozygous for *V. arizonica* specific alleles. As expected, this analysis revealed that portions of chromosome 14 were heterozygous across a region that encompassed Peak 5, Peak 6 (*PdR1*) and beyond (Fig. S12). However, the proximal peak (peak 4) on chromosome 14 was also heterozygous (spanning from ~6.43Mbp to 6.59 Mbp on chromosome 14; Fig. S12). Another prominent peak of heterozygosity on chromosome 9 did not correspond to peaks detected in our GWA. In short, both kmer and SNP analyses suggest that resistance backcrossed from *V. arizonica* encompassed multiple genomic regions.

**Fig. 4.**
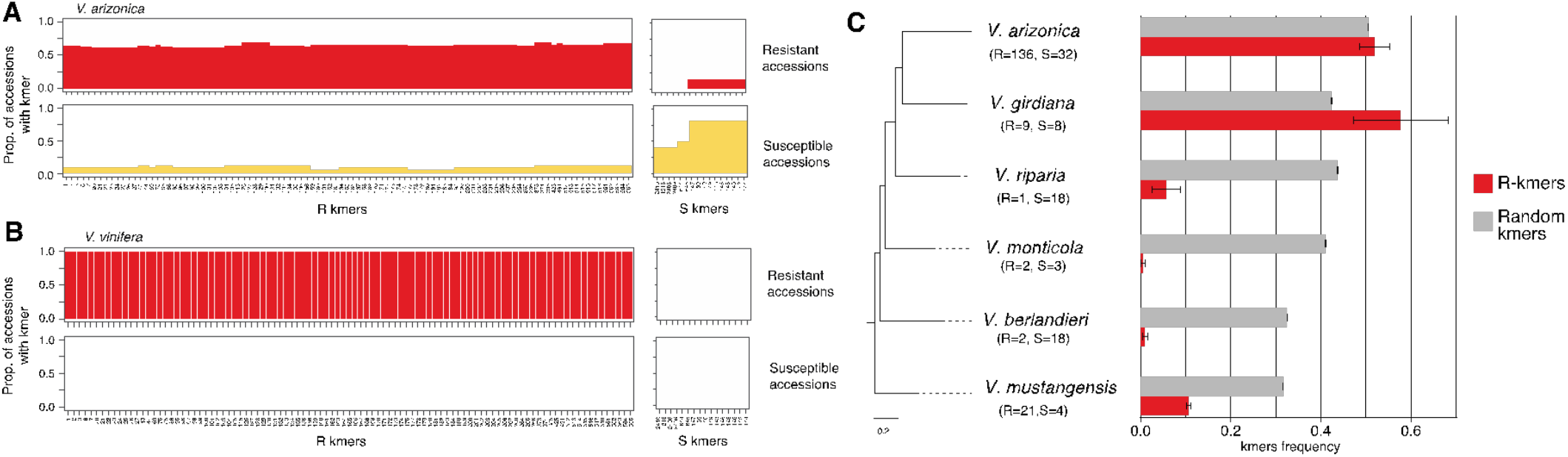
The presence of resistance and susceptibility kmers in different data sets. **A.** Analyses within the *V. arizonica* sample set. The top-left graph indicates the 99 different resistance (R-kmers) kmers across the x-axis, with their detection frequency across the resistant (CFU/mL < 13) accessions. The top-right graph plots the average detection frequency of susceptibility kmers (S-kmers). The bottom-left and bottom-right graph are similar, but show R-kmer and S-kmer detection frequencies among susceptible accessions. **B.** The same graphs as in B, but the top graphs plot R-kmer and S-kmer detection frequencies for the five *V. vinifera* cultivars bred for PD resistance by backcrossing to *V arizonica*, while the bottom graphs represent susceptible *V. vinifera* cutlivars. **C.** Plots of kmer frequencies in six *Vitis* species. The species phylogeny is shown on the left, with the average detection frequency of R-kmers shown in red. The gray bars represent detection frequencies of randomly chosen kmers that had similar population frequencies in *V. arizonica* as the set of R kmers. Whiskers denote standard deviations.

### Resistance markers across *Vitis* species

These observations raise additional questions about the evolution of PD resistance: Did PD resistance arise only once in wild *Vitis* and, if so, is there evidence for the involvement of multiple genic regions? These questions are especially pertinent because all North American wild *Vitis* species can hybridize, with their genomes containing relics of introgression events that are enriched for RLK and RLP genes (Morales-Cruz et al. 2021). To address these questions, we extended kmer analyses to a population genomic sample of 105 individuals from six wild North American *Vitis*, all of which were assayed for resistance (Riaz et al. 2020; Morales-Cruz et al. 2021) (Fig. 4C, Table S8 & Fig. S13). The six species were estimated to have a common ancestor ~25 million years ago (mya) (Morales-Cruz et al. 2021).

We hypothesized that PD resistance was introgressed across species and therefore predicted that the same R-kmers were present across species. We found (as expected) that R-kmers were at significantly higher frequencies in a subset of resistant vs. susceptible individuals for V. *arizonica* (Welch Two Sample WTS t-test, p = 4.50e-16), and also for its sister species, V. *girdiana* (p = 0.007) (Fig. S14). Interestingly, five of the R-kmers within *V. girdiana* mapped to the chromosome 15 peak, again suggesting a multigenic component to resistance. These five kmers were detected in ~67% (12/18) of the *V. girdiana* individuals. These data suggests V. arizonica and V. girdiana share the basis for resistance, with due to introgression or (more parsimoniously) common ancestry. For the remaining four species, no resistant individuals had > 50% of R-kmers (Fig. 4C), with no difference in R-kmer frequency between resistant and susceptible accessions (Fig. S14). In fact, we detected R-kmers less often in these species than for a set of random *V. arizonica* kmers chosen to have similar population genetic frequencies as the R-kmers. Contrary to our hypothesis, the R-kmer distribution in these more distant species provide no evidence that the genetic mechanism of PD resistance (or at least the kmers linked to resistance) was introgressed from *V. arizonica/V. girdiana* to the remaining four species.

### Predicting PD resistance with climate data

Because our plant accessions were sampled across a geographic range (Fig 1), we can use the resequencing data to investigate relationships to climate. We utilized gradient forest (GF) to detect bioclimatic factors related to resistance. GF is a machine learning method that models the turnover in genetic composition and frequency across the climate landscape (Fitzpatrick and Keller 2015) while identifying bioclimatic variables that are important to the construction of the model. As is common for GF applications (Jonás A. Aguirre-Liguori et al. 2021), we applied it to candidate SNPs, specifically the 25 SNPs associated with resistance. To test for robustness, we also repeated GF analysis 1000 times. In all 1000 runs, GF identified BIO8 (Mean Temperature of Wettest Quarter) as the most important model contributor among 10 bioclimatic variables, followed by BIO3 (Isothermality), BIO4 (Temperature Seasonality) and BIO17 (Precipitation of Driest Quarter) (Fig. 5A). Moreover, the turnover function revealed a bias in which susceptible individuals were from locations where BIO8 was <10°C (Fig. 5B), which was confirmed by a significant pairwise comparison between resistant and susceptible individuals (Fig. S15).

**Fig. 5.**
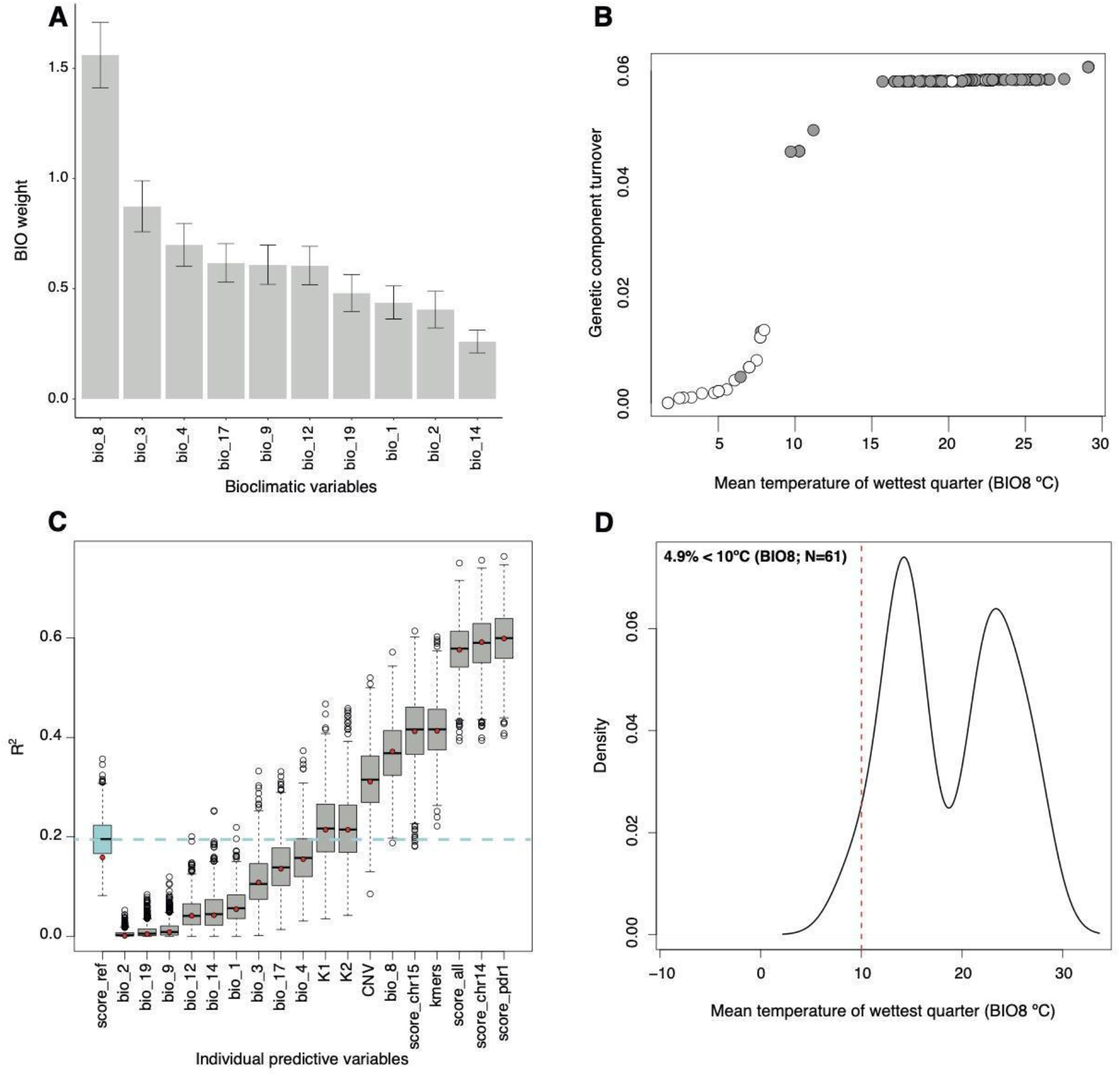
Relationships among resistance, genetic markers and bioclimatic data. **A.** The estimated importance, from GF modeling, of each of the bioclimatic variables tested. The y-axis is a measure of the importance. The histograms denote the average inferred importance of the bioclimatic variable, with the whiskers plotting the standard deviation of 1000 separate analyses. BIO8 was estimated to have the biggest impact on the model in all 1000 analyses. **B**. The turnover function showing the temperature range of BIO8 on the x-axis and the change in the genetic composition on the *y*-axis. The circles represent individuals that are colored by resistance (gray) or susceptible (white). **C**. Individual predictors in a linear model to predict resistance levels (CFU/ml). The label score_ref represents sets of 1000 randomly chosen sets of 25 SNPs. The other sets of predictors include bioclimatic variables and genomic data, as listed in the text, each evaluated 1,000 times with bootstrapped datasets. Each boxplot reports the minimum and maximum values in the whiskers, the quartiles and median values in the square, with the circles showing outliers. The dashed horizontal line reflects the median value of 1,000 replicates of the *R_pd_* score. **D.** The density distribution of BIO8 for a global database of locations of *Xylella fastidiosa* detection.

We performed two additional analyses to assess the generality between resistance and BIO8. First, we examined our complete dataset of all assayed *Vitis* individuals (*n*=275) across six species. The dataset was highly skewed, because 30% of susceptible individuals – but only 1.6% of resistant individuals – had BIO8 < 10°C (FET, p = 8.2e-12). This result held separately for *V. arizonica* (FET, p = 8.2e-12) and when *V. arizonica* was not included in the analysis (FET, p = 0.03). Second, we constructed a global dataset of known *X. fastidiosa* geographic locations that integrates across plant families and all *X. fastidiosa* subspecies (European Food Safety Authority (EFSA) 2018). Unfortunately, the dataset had few exact locations suitable for analysis, leaving only 61 reputable observations. Of these, fewer than 5% had BIO8 values < 10°C (Fig. S15), reflecting the previously reported relationship between temperature and *X. fastidiosa* presence (Purcell 1980; Lieth et al. 2011; Bosso et al. 2016; Sicard et al. 2018).

Given an association between plant resistance and temperature, we explored whether genetic or climatic factors better predicted bacterial load in *V. arizonica*. We assessed individual predictors with linear models, focusing on 10 bioclimatic predictors and nine genomic predictors (Fig. 5**C**). The genomic predictors included kmers, CNVs, assignments into genetic groups (K1 and K2), randomly chosen SNPs, and SNPs associated with PD. We summarized SNPs associated with PD with *S_pd_*, a measure that ranged from 0.0 to 1.0 and reflected the average proportion of alleles associated with resistance (where 0.0 is no resistance-associated alleles).

Focusing on the performance of each of the 19 predictors, *S_pd_* calculated across *PdR1* SNPs (10 total SNPs) had the strongest predictive power (R^2^ = 0.599), followed by related *S_pd_* scores based on all candidate SNPs on chromosome 14 (16 SNPs, R^2^ = 0.592), all candidate SNPs across the genome (25 SNPs, R^2^ = 0.576) and finally all candidate SNPs on chromosome 15 (6 SNPs, R^2^ = 0.412) (Fig. 5C). Among the bioclimatic variables, BIO8 had an R^2^ of 0.370 in the linear model, which was much higher than the median value for 1000 randomly chosen sets of SNPs (R^2^ =0.196) and similar to the predictive power of Kmers (R^2^ = 0.410) and CNVs (R^2^ =0.307). Thus, BIO8 was a reasonable predictor of resistance, even in the absence of genetic data. Notably, the other bioclimatic variables that were identified by GF were not strongly predictive by themselves - e.g., BIO4, BIO17 and BIO3 had lower predictive power than random sets of SNPs (Fig. 5C).

## DISCUSSION

*X. fastidiosa* causes Pierce’s Disease in domesticated grapevines (*V. vinifera*) and economically devastating diseases in other crops like citrus, coffee and almonds (Rapicavoli et al. 2018). A diverse body of work has investigated the basis of resistance across diverse crop species but few plausible candidate resistance genes (Rodrigues et al. 2013; Giampetruzzi et al. 2016; Zaini et al. 2018; Agüero et al. 2022). To date, however, no studies of *X. fastidiosa* resistance have taken advantage of full-scale genomic approaches like GWA. Indeed GWA studies in the wild relatives of crops are surprisingly rare, despite the importance of understanding the basis of resistance in ecological settings (Bartoli and Roux 2017) and the transformative potential of such knowledge for crop breeding (Migicovsky and Myles 2017). Here we have applied GWA to resistance in *V. arizonica*, based on an improved reference genome, on resequencing data from 167 wild-sampled accessions and on phenotypic data measured from *X. fastidiosa* infection assays (Riaz et al. 2020; Morales-Cruz et al. 2021). Together, these analyses have yielded information about genomic regions associated with PD resistance and also identified candidate genes within those regions. Moreover, the landscape scale of the resequencing data has allowed us to explore relationships between plant resistance and climate.

### Genome-wide associations with PD resistance

Not surprisingly, GWA identified several significant SNP and kmer markers on chromosome 14 between the genetic boundaries that define the *PdR1* locus (Fig. 2). This locus was originally identified by genetic mapping and QTL analyses (Doucleff et al. 2004; Krivanek et al. 2006) and it was subsequently backcrossed into susceptible *V. vinifera* to produce resistant cultivars (Riaz et al. 2009). There had been no insights into causative genes that lie within this region, until Aguero et al. (2022) (Agüero et al. 2022) identified five putative R genes and transformed two LRR genes separately into *V. vinifera*, failing to find evidence of enhanced resistance. Our study of a different accession (b40-14) provides further insights. First, based on careful genomic annotation combined with association and gene expression analyses, we have identified a narrower *PdR1* region with two strong candidates (g238150 and g238180), along with two additional candidates (g238250 and g238290) within the traditional *PdR1* locus (**Fig. 3**). Second, we have mapped the two transformed LRR genes from Aguero et al. (2022) to our genome (see Methods); neither of the two transformed genes are closest homologs to candidate genes in b40-14 and neither are highly expressed in b40-14 (Table S9). Overall, our candidate genes, which are present on both b40-14 haplotypes, differ substantially from those identified in b43-17. It is possible, of course, that different genes confer resistance in different accessions, given that structural variants are common in *Vitis* genomes (Zhou et al. 2019) and that allelic heterogeneity for resistance is also common (Karasov et al. 2020). A reasonable next step is to knock-out our candidates in *V. arizonica*, but unfortunately transformation in *Vitis* is currently efficient primarily for a narrow set of *V. vinifera* cultivars (Zhang et al. 2021). An important challenge for viticulture is to improve transformation techniques for application to more cultivars and to agronomically valuable wild species.

One interesting possibility is that candidate genes within *PdR1* are not sufficient to confer resistance. There is evidence consistent with and against this hypothesis. Some of the evidence is historical. In an early study of PD resistance among *Vitis* species, Mortensen (1968) performed controlled hybrid crosses and concluded that complementary gene action among three independent genes best explained his results (Mortensen). Our data also hint at the contribution of multiple genomic regions. For example, the striking LD pattern on chromosome 14 suggests additive or epistatic interactions between distinct regions on the chromosome (Fig 3A). Similarly, *V. girdiana* and *V. arizonica* share R-kmers that map to at least two different chromosomes, further reflecting multi-genic composition (Fig. 4B). Finally, genomic investigation of resistant *V. vinifera* cultivars demonstrate that they include backcrossed contributions from *V. arizonica* across the *PdR1* region, as expected, but also across additional peaks on chromosomes 8, 9, 14 and 15 (Fig. 4A). In contrast to this multigenic interpretation, we have observed that most significantly associated markers lie within the *PdR1* region (Fig 2) and also that resistance can be best predicted using SNP data from within *PdR1* (Fig. 5B). We have not estimated the effect sizes of SNP and kmer markers directly, because the gold standard for effect estimation requires a second independent dataset (Burghardt et al. 2017). However, our prediction results are consistent with the loci of largest effect residing within *PdR1*. Finally, a QTL study based on another *V. arizonica* parent found that 55.5% of phenotypic variation of resistance is explained by markers on chromosome 14 (Huerta-Acosta and Riaz), consistent with a major but not complete *PdR1* effect. Overall, we interpret these complex observations to predict: *i*) that a combination of the correct candidate genes within *PdR1* may confer substantial resistance but *ii*) that resistance at levels mimicking those in the wild will require allelic variants from additional genomic regions.

### Evolutionary and ecological insights into plant resistance

Inferring the spatial distribution of disease resistance is critical for understanding its evolution and ecology (Karasov et al. 2020). We have investigated resistance across the landscape of *V. arizonica* (Fig. 1) and across six wild species that segregate for PD resistance. Given that all North American *Vitis* species are interfertile and that there is ample genomic evidence for historical introgression of resistance genes among species (Morales-Cruz et al. 2021), we predicted that the genetic solution to PD would yield clear signals of introgression. We find that R-kmers are commonly shared between *V. arizonica* and its closest species in our sample, *V. girdiana*, but not across the other species (Fig. 4C). It is possible that we cannot detect introgression events because causative loci and associated R-kmers have become uncoupled over evolutionary time. At a minimum, however, our results provide no evidence that PD resistance has introgressed from *V. arizonica* to the non-*girdiana* species in our sample. If true, this implies that other wild *Vitis* species have independently evolved mechanisms of PD resistance; hence further study of these species may provide additional insights into alternative genetic causes of PD resistance. In this vein, one particularly interesting species is *V. mustangensis* (syn *V. candicans*), which has not been widely utilized agronomically as a rootstock or for hybrid scion breeding, but it does segregate for PD resistance and also contains alleles that may be useful for viticulture in the context of climate change (J. A. Aguirre-Liguori et al. 2021).

We have taken advantage of our geographic sampling to investigate correlates between resistance and climate, finding that resistance-associated SNP markers correlate with climatic variables, especially temperature in the wettest quarter (BIO8). More specifically, susceptible plants tend to be found where BIO8 is <10°C; this is true within *V. arizonica*, within our expanded sample of *Vitis* species and across wider (although still quite limited) geographic samples of *X. fastidiosa* that integrate across plant hosts and bacterial subspecies (European Food Safety Authority (EFSA) 2018) (Fig 5D). Somewhat remarkably, this simple climatic measure predicts bacterial load nearly as well as some genetic markers (kmers) and better than others (CNVs) (Fig 5D). This is, to our knowledge, the first time that genomic data have been used to associate plant resistance and climate, yielding a useful bioclimatic predictor. Our findings are not without precedent, however, because temperature has previously been identified as a strong predictor of *X. fastidiosa* distribution and presence (Purcell 1980; Lieth et al. 2011; Sicard et al. 2018; Raffini et al. 2020). Combining these observations, we suspect that individuals with low (<10°C) BIO8 temperatures lack resistance because *X. fastidiosa* growth is hampered at low temperatures (Hopkins and Purcell 2002; Bosso et al. 2016) and/or because temperature affects its insect vectors (Hoddle 2004; M. Godefroid et al. 2022). Plant resistance will not be favored by natural selection in regions where the pathogen does not persist, particularly if there is a fitness cost to resistance [as has been demonstrated for R-gene mediated resistance in *A. thaliana* (Tian et al. 2003)].

Previous work has modeled the distribution of *X. fastidiosa* under climate change (Purcell 1980; Lieth et al. 2011; Sicard et al. 2018; Raffini et al. 2020; Martin Godefroid et al. 2022), but these models have depended primarily on incomplete *X. fastidiosa* surveys and have not been informed by data on the distribution of plant resistance. To illustrate how such information may be useful, we have employed climate models to predict where BIO8 will shift in the future. More specifically, we have identified regions where BIO8 will transition across the threshold of BIO8 =10°C (Fig. 6A), as informed by plant resistance. By categorizing regions where BIO8 is predicted to move from below (or above) 10°C in the present to above (or below) 10°C in the future and by assuming the distribution of resistance informs on pathogen presence, we can identify regions where *X. fastidiosa* pressure is likely to expand or contract. We performed these categorizations across 54 climate models to consider uncertainty in global circulation models, shifts in greenhouse gas emissions and time (see Methods). Summarizing across models, we have found that most of the globe will not transition - i.e., it will remain either above or below 10°C during the wettest quarters (Fig. 6A). Perhaps unsurprisingly (Cohen et al. 2021), large portions of Canada, Eastern Europe, Russia and Northern Asia climate are predicted to move beneath the BIO8 threshold, suggesting that these regions may be less likely to have pathogen pressure in the future. There are, however, discrete areas of the Western Americas, Western Europe, Central Asia, Southern Australia and elsewhere that are predicted to transition above the 10°C threshold in most models, suggesting increasing *X. fastidiosa* pressure in these regions (Fig. 6A). These models of course make numerous assumptions. Some are common to all climate models (e.g., a reliance on accuracy of the climate predictions) and to previous *X. fastidiosa* species distribution models [e.g., which ignore the potential of *X. fastidiosa* to evolve to new temperature regimes (Jonás A. Aguirre-Liguori et al. 2021; Martin Godefroid et al. 2022)]. Some are more specific to this work - i.e., that *X. fastidiosa* is not dispersal limited and also that the BIO8 threshold has importance beyond *V. arizonica*, as suggested by our analysis of data that includes different plant hosts (Fig. 5C). Importantly, however, our climate modeling illustrates how data about plant resistance can help inform the potential distribution of disease under a shifting climate.

**Fig. 6.**
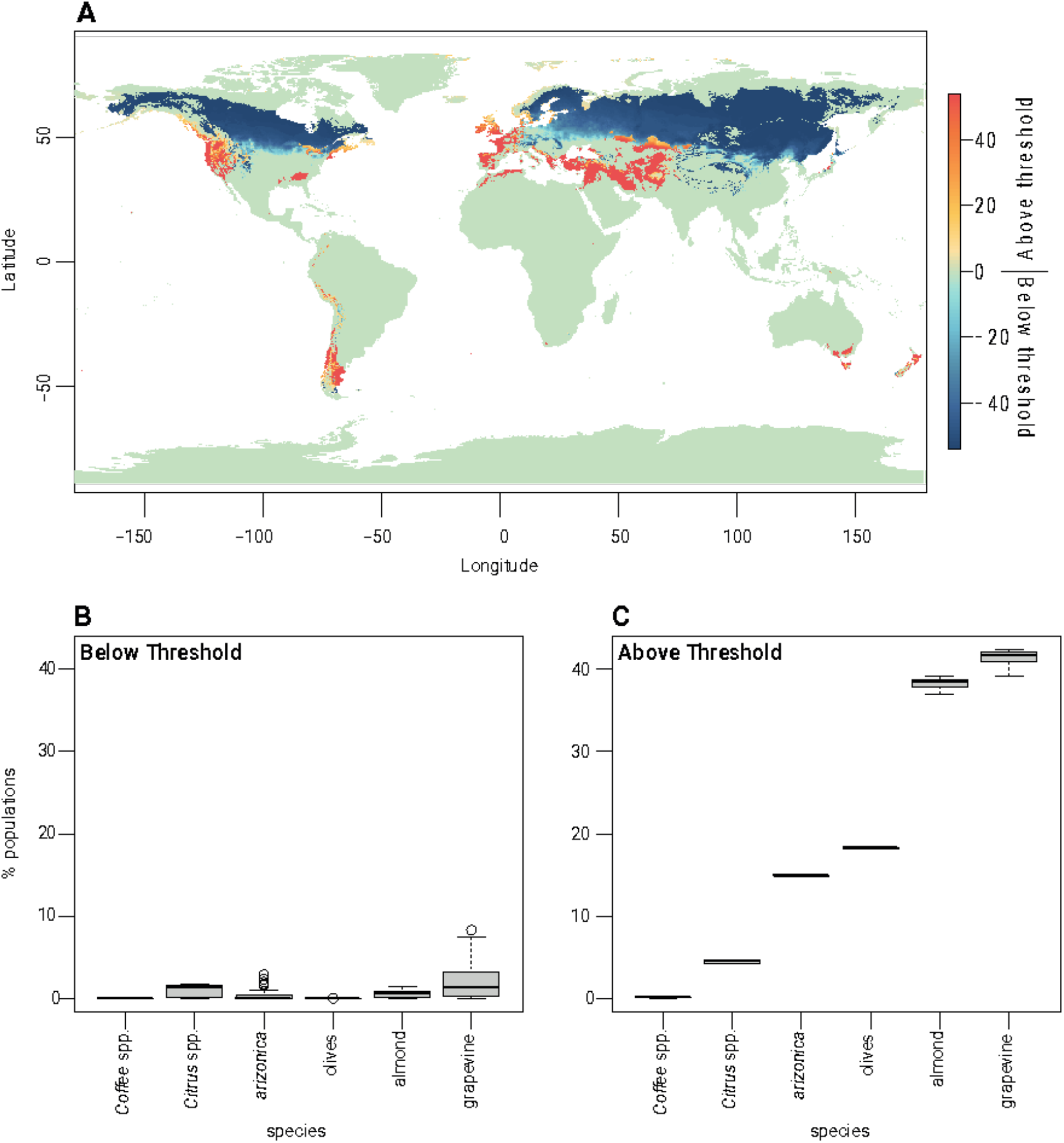
Climate predictions and projections of the prevalence of Xylella fastidiosa for focal crops. **A**. The map portrays the number of climate models (out of 54 total) that support movement across the BIO8 = 10°C threshold. The warmer colors reflect regions that are moving from below (in the present) to above the threshold, while the cooler colors portray ares that that are moving from above (in the present) to below the threshold. **B**. A summary of the percentage of locations associated with movement from above the 10°C threshold (in the present) to below the threshold for five crops and *V. arizonica*. **C**. A summary of the percentage of locations associated with movement from above the 10°C threshold (in the present) to below the threshold for five crops and *V. arizonica*. Both B and C are based on 6,204 locations for coffee; 3,386 locations for almonds, 1,111 locations for V. arizonica; 5, 256 locations for Citrus species; 174,713 locations for olives and 33,225 location for grapevines.

In fact, we can proceed one step further by assessing the potential effects of climate on specific plant taxa - i.e., wild *V. arizonica* and five affected crops (grapes, coffee, almonds, citrus and olives). To do so, we first downloaded the global locations where each species is grown and then used climate projections to estimate the proportion of locations that will transition over the 10°C BIO8 threshold under climate models (see Methods). Using this approach, we predict that few locations for these crops will transition below the 10°C threshold. In citrus and grapes, for example, the average estimate across the 54 climate models is that only 0.97% (of 7,853) locations and 2.10% (of 40,075) locations will transition below 10°C under climate models (Fig. 6B). Perhaps not unexpectedly, a much higher proportion of locations are estimated to exceed the 10°C threshold over time (Fig. 6C). For example, we estimate that >38% of regions of almond and grapevine cultivation will transition above the 10°C threshold thus, given our assumptions, providing *X. fastidiosa* more favorable conditions and hence potentially subjecting these crops to higher pathogen pressure. Similarly, 18% of olive growing locations are expected to transition above the 10°C threshold, which adds future concern to models that strongly predict the spread of the bacterium to olive growing regions under current climates (Schneider et al. 2020). These calculations treat *X. fastidiosa* subspecies similarly, even though there are some differences in their host specificity (Sicard et al. 2018; Batarseh et al. 2022) and distributions (Godefroid et al. 2019; Schneider et al. 2020), but emphasize that climate change is unlikely to affect all crops similarly with respect to *X. fastidiosa* exposure.

Overall, our study has considered the genomic, climate and evolutionary context of resistance to a pathogen that is an emerging global pest and that already causes devastating economic damage to several crops. By studying the complex genetic architecture of PD resistance in a wild grapevine, we have implicated several genomic regions in resistance and identified genes that are fitting candidates for genetic introduction into susceptible crops. Furthermore, by comparing features across wild *Vitis* species we have uncovered hints about the origins of this critical trait. Finally, our work has highlighted the potential of combining landscape-scale resequencing data and climate models to predict the shifting pressures of a damaging plant pathogen. These results underscore the urgent need to identify additional *X. fastidiosa* management and containment methods, potentially via enhanced information about resistance mechanisms and genes.

## MATERIALS AND METHODS

### Plant material and PD disease evaluations

We studied 167 accessions of *V. arizonica* collected across the southwestern states of the US and northern Mexico, covering the distribution of the species (Figure 1A and Fig. S1). The sampling location was available for all 167 *V. arizonica* accessions, which are part of a living collection of Southwest *Vitis* accessions maintained at the University of California, Davis; all *Vitis* accessions used in this paper were from that collection. PD resistance in these 167 accessions were previously assayed in controlled greenhouse trials (Riaz et al. 2020; Morales-Cruz et al. 2021), following previously published protocols (Krivanek and Walker 2005; Riaz et al. 2020). Briefly, accessions were inoculated with *Xylella fastidiosa*, and the susceptible *V. vinifera* cultivar Chardonnay as a reference control. Nineteen screens were carried out from 2011 to 2020, with a minimum of four biological replicates per accession. Disease severity was assessed 10 to 14 weeks after inoculation using categorical phenotypes for disease severity, and ELISA was used to measure bacterial load (colonizing forming units or CFUs/ml) in the stem above the inoculation site from 12 to 14 weeks. The ELISA data were log-transformed and statistical analysis was performed using JMP Pro14 software (Copyright 2020, SAS Institute Inc.) to determine the variability of ELISA for the reference control plants across experiments. In this study, we used the quantitative measure of plant bacterial load (Least Squared Means of colonizing forming units or CFUs/ml) as an indicator of disease resistance. Lower bacterial load reflects higher PD resistance.

### Genomic reference, resequencing data and SNP calling

Version 1 of the *V. arizonica* sequence (b40-14 v1) was published previously (Massonnet et al. 2020; Morales-Cruz et al. 2021), but this study relied on an updated version (b40-14 v2). Version 2 was a complete re-assembly based on the application of Haplosync (Minio et al. 2022) combined with ~ 2000 rhAmpSeq Vitis markers (Zou et al. 2020), independent of any single reference genome assembly. This new version of *V. arizonica* genome chromosome haplotypes are better phased, with fewer unplaced sequences (~ 65 Mb; 3,035 gene loci). The correct phasing of the locus was assessed during the assembly quality control steps based on genome and locus-specific markers, and gene content. The gene annotation was ported between versions 1 and 2. The genome now contains 57,003 gene loci, a number slightly changed due to gene model corrections and homozygous filling of the haplotypes during the assembly. The version 2 assembly represents the most contiguous genome assembly of any wild *Vitis* genome released to date and was used as the reference for all analyses. The genome is available for browsing at *grapegenomics.com*, but the v2. assembly and fasta files are also available from #### (will be freely available before publication).

Our *V. arizonica* resequencing dataset consisted of short-read, whole genome data from 167 *V. arizonica* individuals, for which a subset of *n*=22 had been sequenced previously (Morales-Cruz et al. 2021). The data for this subset was available from NCBI BioProject PRJNA607282. For the remaining 145 individuals, genomic DNA was extracted from leaf samples with the Qiagen DNeasy plant kit. Sequencing libraries were constructed with an insert size of ~300 bp using Illumina library preparation kits and were sequenced using the Illumina HiSeq 2500 platform with 2 × 150 bp paired reads to a target coverage of 10x. The raw sequencing data for this study has been deposited in the Short Read Archive at NCBI under BioProject ID: PRJNA842753.

We filtered and evaluated raw reads from 167 individuals using Trimmomatic-0.36 (Bolger et al. 2014) and FastQC (Andrews and Others 2010). Filtered reads were then mapped to the reference genome (independently to Hap1 and Hap2) with the BWA-MEM algorithm (Li 2013) implemented in bwa-0.78. Joint SNP calling was conducted using the GATK v.4.2.2.0 pipeline (McKenna et al. 2010) for Hap1 and Hap2 independently. We first used the integrated version of Picard tools (Institute 2016) to remove duplicated reads with the “MarkDuplicates” function, followed by the “AddOrReplaceReadGroups” function to label the reads for each individual. For the SNP prediction we used the HaplotypeCaller algorithm with a sample ploidy of 2 and a mapping base quality score threshold of 20 (Q > 20). We combined the VCF files of all individuals to make the final SNP calls using the “GenotypeGVCFs” function with default parameters. We then filtered raw SNPs with bcftools v1.9 (Li et al. 2009) (https://samtools.github.io/bcftools/) and vcftools v0.1.17 (https://vcftools.github.io/) (Danecek et al. 2011). We kept SNP sites for downstream analysis if they were biallelic, had quality higher than 30, had a depth of coverage higher than five reads, had no more than three times the median coverage depth across accessions, and had no missing data among individuals. Additionally, the following expression was applied under the exclusion argument of the filter function in bcftools: “QD < 2.0 | FS > 60.0 | MQ < 40.0 | MQRankSum < −12.5 | ReadPosRankSum < −8.0 | SOR > 3.0”.

### Population Structure and genome-wide associations

We transformed VCF the file into BEAGLE format using vcftools 0.1.17. We used BEAGLE files as input to evaluate the genetic structure of the *V. arizonica* using the NGSAdmix software included in the ANGSD package version 0.931-21-g13af014 (Korneliussen et al. 2014). We ran NGSadmix for 1 to 10 ancestral populations (*K*), repeating analyses 10 times for each *K* value and including variants with a minimum minor allele frequency > 0.05 (Fig. S16). We then employed the Cluster Markov Packager Across K (Clumpak) software (Kopelman et al. 2015) to detect the *K* value with the highest likelihood of *K* = 2.

The NGSadmix results were used to help guide controlling for genetic structure in genome-wide association (GWA). We performed GWA to identify significant associations between SNP allelic frequencies and bacterial load (*Xylella fastidiosa* CFU’s), using two different methods that control for population structure using different approximations. First, we used LFMM2, which uses latent factors to control for genetic structure (Frichot et al. 2013). The number of latent factors were chosen based on a visual observation of the screenplot of the percentage of variance explained by the loadings of the genetic PCA of all individuals. The PCA was obtained using all loci with no missing data and with the prcomp function in R (Kassambara). In total, we defined K=4 latent factors. Next, we ran LFMM2 using a ridge penalty (function lfmm_ridge), and we controlled for genomic inflation factor (function lfmm_test, with calibrate=“gif”). We corroborated that the *p* values had a flat distribution, and we corroborated that the genetic structure was well controlled based on a qqplot (Fig. S4 & S7).

In addition, we used the variance component algorithm Efficient Mixed-Model Association eXpedited (EMMAX) version beta-07Mar2010 (Kang et al. 2010). We converted the set of filtered SNPs with no missing data from VCF format to transposed ped format using PLINK version 1.90b6.16 (Purcell et al. 2007). Using the transposed ped files as input, we calculated the Balding-Nichols (BN) kinship matrix using the “emmax-kin” script and default parameters. Finally, we ran the associations using the SNPs data (as transposed ped), the BN matrix, and the phenotype as Least Squares Means of CFU/ml of *X. fastidiosa* for each accession. Both methods (EMMAX and LFMM2) were adjusted for multiple comparisons using the Bonferroni correction of the program “p.adjust” from the stats package version 4.1.2 in R. We focused only on SNPs and candidate regions that were detected by both methods.

### Kmer-based GWA

To perform GWA based on kmers, we followed a previously published pipeline (Voichek and Weigel 2020) (https://github.com/voichek/kmersGWAS). Briefly, we extracted all kmers and canonized (i.e. reverse complement is assumed to be the same kmer) kmers of 31 bp in size using KMC version 3 (Kokot et al. 2017). We extracted the kmers directly from the paired and unpaired filtered reads for each of the 167 *V. arizonica* samples independently. We compared the kmers across samples and created a table of kmers that were found in at least 5 individuals (“-mac 5”) and in each canonized/non-canonized form in at least 20% of individuals from which it appeared in (“-p 0.2”). We used the script “emma_kinship_kmers” included in the pipeline with a MAF < 0.05 filter to create a kinship matrix based on the kmer table. Finally, we ran the kmer-GWAS with the script “kmers_gwas.py” and GEMMA version 0.98.5 (Zhou and Stephens 2012) with the kinship matrix, the kmer table, and the phenotype data as Least Squares Means of CFU/ml of *X. fastidiosa* for each accession. The script provided a list of 9991 kmers that passed a parametric test as an initial filter. To filter more stringently, we used the number provided in the file “pheno.tested_kmers”, which was 967066440, to adjust the *p*-values with a Bonferroni correction using the program “p.adjust” from the stats package version 4.1.2 in R. To create a textual version of the presence/absence kmers of the significant Kmers we used the “filter_kmers” from the pipeline. Given 115 significant kmers, we mapped them to the *V. arizonica* genome using BLASTN (Zhang and Madden 1997) and a word size of 8 bp. We filtered all the BLASTN results and kept alignments, allowing a maximum of 1 mismatch.

We further explored the sequences of the 36 kmers that did not map to our reference genome. We first searched and extracted the reads matching these 36 kmers across the 167 *V. arizonica* accessions. We then used SPADES v3.15.4 (Bankevich et al. 2012) to assemble the matching reads into contigs independently for each kmer. Finally, we aligned the resulting contigs to the Reference RNA Sequences database (Refseq_rna) using BLASTN (visited on 06/10/2022) and recorded the top hit gene.

### Copy Number variation analysis and associations

To identify Copy Number Variants (CNVs), we used the program CNVcaller version 2.0 (https://github.com/JiangYuLab/CNVcaller) (Wang et al. 2017). CNVcaller uses normalized read-depth values across windows in the genome to identify CNVs and it is especially suited for large population data like our *V. arizonica* sample. First, we generated a duplicated window record file specifically from our genome reference *V. arizonica* b40-14 v2 using a window size of 2 kb and for each chromosome independently. We then analyzed the individual read depth in 2 kb non-overlapping windows using the “Individual.Process.sh” script and the alignment files of all 167 accessions in bam format with the PCR duplicated reads removed during the SNP calling pipeline (see section above). The script produces normalized values of read depth across the genome for each genotype. We then used the normalized read depth values of all genotypes as input to the script “CNV.Discovery.sh”, excluding windows with a lower frequency of gain/loss individuals of 0.1 (“-f 0.1”) and with Pearson’s correlation coefficient lower than 0.3 (“-r 0.3”). Finally, we used “Genotype.py”to classify genotypes across the population according to their CNV profiles.

To explore the associations between CNVs and PD-resistance, we used the CNVcaller estimation of diploid copy number for each CNV and tested for correlations with *X. fastidiosa* bacterial levels, while taking into account the genetic structure of the *V. arizonica* population. We used the R library “ppcor” v1.1 (Kim 2015) to run a partial correlation for each of the CNVs, using genotype assignment (*Qi*) values from the genetic structure analysis (see section above) as the confounding variable. To account for multiple testing we imposed a Bonferroni correction and identified significant CNVs with adjusted p< 0.05.

### Defining PD-associated peaks

We performed a total of four association analyses: LFMM2 based on SNPs, EMMAX based on SNPs, kmer-based GWA, and CNV-based GWA. From these analyses we defined eight GWA peaks of interest in the genome (Figure 2). To define these peaks, we required that a peak contains at least one SNP that was significant with both LFMM2 and EMMAX. However, most peaks had multiple pieces of evidence - i.e., either more than one SNPs, significant kmers and/or CNV variants. When applying this logic, we focused only on kmers that mapped uniquely to the genome and so excluded 17 kmers that mapped to multiple places in the genome with the same identity and alignment length.

### Analyses of PD-associated kmers in other *Vitis* species

We identified kmers associated with resistance in *V. arizonica* sample and then characterized their presence in three different resequencing datasets: *i*) a multiple species *Vitis* dataset, *ii*) a dataset generated from scion cultivars bred for PD resistance by backcrossing to the b43-17 accession of *V. arizonica*, and *iii*) a set of PD susceptible cultivars. The first dataset included 105 accessions from five species: *V. arizonica* (n =22), *V. candicans* (n = 24), *V. berlandieri* (n = 22), *V. girdiana* (n = 18) and *V. riparia* (n = 19) (Morales-Cruz et al. 2021). All of these accessions had been assayed for PD resistance, and categorized as resistant if CFU/mls were <13.0 at the time of assay. In this dataset, 20 *V. arizonica* were resistant (n=20), 21 *V. candicans* (n = 25), 3 *V. berlandieri* (n = 21), 9 *V. girdiana* (n = 17), 2 *V. monticola* (n=5) and 2 *V. riparia* (n = 20). The resistance assay data and sampling locations of the accessions are available (Morales-Cruz et al. 2021). For the second dataset, we generated resequencing data for five PD resistant cultivars (Ambulo Blanc, Caminante Blanc, Camminare noir, Errante noir and Paseante noir). DNA extraction, library preparation and Illumina sequencing followed the protocols mentioned above, and the data were deposited into Bioproject: PRJNA842753. Finally, the ‘control’ dataset of PD susceptible accessions was downloaded from public databases (Cabernet Sauvignon cl. 08: SRR3346862; Chardonnay cl. 04: SRR5627799; Zinfandel cl. 03: SRR8727823; and Petite Sirah: SRR12328988). For each of the datasets, we generated kmers of 31 bp for each sample as described above. We then searched for the presence of 115 associated kmers from *V. arizonica* using the “filter_kmers” script from the kmer-GWAS pipeline (Voichek and Weigel 2020) (https://github.com/voichek/kmersGWAS).

To compare the R-kmers with random sequences in other *Vitis* species, we first extracted 100,000 unique and random kmers from the *V. arizonica* population. We then calculated the frequency of these sequences across all individuals and selected kmers with similar frequencies as the R-kmers mean (0.52), resulting in 38,523 kmers. We then created 100 subsets of 99 random kmers from the set with similar frequencies as R-kmers. Finally, we searched for the presence of each set of other *Vitis* species and calculated the mean frequencies of the 99 kmers in each random set. We report the average and standard error of means across the 100 sets in Fig. 4C.

### Linkage disequilibrium

We calculated the genome-wide LD decay across the *V. arizonica* population with the software PopLDdecay v3.40 (Zhang et al. 2019). We used the filtered SNPs of the 167 individuals from hap1, allowing a maximum distance of 1 Mb (Fig. S11). We used the perl script “Plot_OnePop.pl” included in the package to create the decay graph.

To explore the LD landscape of the regions around PdR1 and chromosome 14 as a whole we used Tomahawk v0.7.0 (https://github.com/mklarqvist/tomahawk). We used as input the filtered SNPs of chromosome 14 for hap1 as input, containing the 167 *V. arizonica* accessions. We converted the VCF file into a custom format file (“.twk”) for the package and calculated the LD with the “calc” function. We then filtered LD values using the “view” function, keeping regions with R^2^ > 0.5 and p-values < 0.001. Given that we were interested in the LD at the chromosome-scale (~30Mb) we used the “aggregate” function. Using this function we aggregated R^2^ values in 1000 bins for both the x and y-axis, and used 5 as the minimum cut-off value in the reduction function. Finally, we used the “rtomahawk” R package (https://github.com/mklarqvist/rtomahawk) to create the chomosome-scale LD landscape plot using the aggregated R^2^ values.

### Functional annotation and Refinement of *PdR1* gene models

Gene models located within the two haplotypes of *Pdr1* were manually refined by visualizing alignments of RNA-seq reads from *V. arizonica* b40-14 leaves (Morales-Cruz et al. 2021) using Integrative Genomics Viewer (IGV) v.2.4.14 (Robinson and Others 2017). RNA-seq reads were aligned onto the diploid genome of *V. arizonica* b40-14 using HISAT2 v.2.1.0 (Kim et al. 2015) and the following settings: --end-to-end --sensitive -k 50.

Predicted proteins of *Pdr1* genes were scanned with hmmsearch from HMMER v.3.3.1 (http://hmmer.org/) and the Pfam-A Hidden Markov Models (HMM) database (El-Gebali et al. 2019) (downloaded on January 29th, 2021). Protein domains with an independent E-value less than 1.0 and an alignment covering at least 50%of the HMM were selected. Transmembrane helices were predicted with TMHMM2 v2.0c (Krogh et al. 2001). Proteins with a predicted Leucine Rich Repeat (LRR) domain and a transmembrane helix were classified as LRR receptorlike proteins. Proteins having a predicted LRR or lysin motif (LysM), a kinase domain, and transmembrane helices were categorized as receptor-like kinases.

Predicted proteins of *Pdr1* genes were aligned onto the predicted proteome of *A. thaliana* and the grape PN40024 (V1 annotation) using BLASTP v.2.2.28+ (Altschul et al. 1990). Alignments with an identity greater than 30% and a reciprocal target:query coverage between 75% and 125% were kept. For each *V. arizonica* protein, best hit in the *A. thaliana* and PN40024 proteomes was determined using the highest product of identity, query coverage, and reference coverage. The sequences of the two ORFs (V.ari-RGA14 and V.ari-RGA18) (Agüero et al. 2022)were aligned onto the b40-14 genome using blastn v. 2.2.28+ (Altschul et al. 1990). For V.ari-RGA14, the alignments had highest coverage of 99.46% and and identity of 100% and located at 26.666 Mb on chromosome 14, which was within the *PdR1* locus but not a strong candidate based expression analyses (Fig. 3BC). For V.ari-RGA18, the alignment with the highest coverage (99.76%) and identity (100%) was located at position 24.283 Mb, well outside the *PdR1* region based on our b40-14 reference (Fig. 3B).

### Gene expression analyses

To evaluating the transcript abundance of candidate genets, plants from three PD-resistant genotypes and three PD-susceptible genotypes of the 07744 population (*[V. rupestris* Wichita refuge x *V. arizonica* b40-14] x *V. vinifera* Airen) were propagated in a controlled environment and inoculated with either *Xylella fastidiosa* or water. Pieces of green stem at 10, 20, 30 and 40 cm above the inoculation were collected from each plant at 1, 2, 3, and 4 weeks post inoculation. Pieces of green stem from each genotype were pooled together. Each genotype constitutes a biological replicate. All plant material was immediately frozen in liquid nitrogen after collection and ground into powder. Total RNA were extracted as described in Massonnet et al. (2017) (Massonnet et al. 2017). cDNA libraries were prepared using the Illumina TruSeq RNA sample preparation kit v.2 (Illumina, CA, USA) and sequenced in single-end 100-bp reads on an Illumina HiSeq4000. RNA-seq reads were parsed using Trimmomatic v.0.3 (Bolger et al. 2014) with the following settings: LEADING:7 TRAILING:7 SLIDINGWINDOW:10:20 MINLEN:36. Transcript abundance was evaluated with Salmon v.1.5.1 (Patro et al. 2017) with the parameters: --gcBias --seqBias --validateMappings. A transcriptome index file was built using a k-mer size of 31 and the combined transcriptomes of *V. arizonica* b40-14 (v2.1), *V. vinifera* cv. Cabernet Sauvignon (Massonnet et al. 2020), with their genomes as decoy. Quantification files were imported using the R package tximport v.1.20.0 (Soneson et al. 2015).

### Bioclimatic variables associated with resistance

To identify the association between SNPs associated with resistance and the environmental landscape, we applied gradient forest (GF) (Ellis et al. 2012), which models the turnover in genetic composition across the landscape (Fitzpatrick and Keller 2015) and identifies both the bioclimatic variables that contribute to the construction of the model and the ‘turnover function’ - i.e., the change of genetic composition across the landscape (Fitzpatrick and Keller 2015; Capblancq et al. 2020; Waldvogel et al. 2020). To estimate the GF model, we used the *gradient forest* package in R, using the 25 SNPs identified by LFMM2 and EMMAX as response variables and using bioclimatic variables as predictive variables. The 19 bioclimatic variables were filtered to retain any correlations < 0.80, based on a variance inflation factor calculated by corSelect from the R package *fuzzySim* (Barbosa 2015). After filtering, we retained 10 of 19 bioclimatic variables (BIO1, BIO2, BIO3, BIO4, BIO8, BIO9, BIO12, BIO14, BIO17, BIO19). We performed the GF analysis using SNP frequencies from each individual (i.e., 0, 1 and 0.5 for heterozygotes) (J. A. Aguirre-Liguori et al. 2021) and repeated the analysis 1,000 times. We also plotted the turnover functions for each bioclimatic variable to show how populations with different resistance to PD are distributed across allele frequency change (Fig. S15).

### Predicting PD resistance

To determine whether genetic and environmental variables predicted PD resistance, we first ran a linear model individually for each bioclimatic and genetic variable to determine their individual predictive power, using the *lm* function in R. We performed 1,000 bootstrap replicates for each model to estimate the variance in predictive ability.

For the bioclimatic variables, we estimated the individual linear models between the 10 bioclimatic values from where the *arizonica* individuals were sampled and their bacterial load.

For some genetic variables, we first estimated a polygenic score across SNPs that associated with bacterial load. This polygenic score, which we called the PD score (*S_pd_*), was calculated as the proportion of alleles that contribute to resistance to PD in a given individual. The state of these contributing alleles was inferred from the GWA results. *S_pd_* was designed after the population adaptive index, which measures the proportion of alleles that show patterns of local adaptation (Bonin et al. 2007). *S_pd_* assumes that alleles contribute equally to PD resistance and that a higher proportion of resistance alleles correlates with lower CFU in the individuals*. S_pd_* ranges from 0 when no resistance alleles are present to 1 when all alleles across loci are homozygous for the resistant state. We estimated *S_pd_* for different sets of SNPs: *i*) all candidate SNPs across the genome (25 SNPs), *ii*) all candidate SNPs on chromosome 14 (16 of the 25 SNPs); *iii*) all candidate SNPs in the region defined by *PdR1* (10 of 25 SNPs); *iv*) all candidate SNPs on chromosome 15 (6 of 25 SNPs). To control for potential ancestry effects, we also tested an analogous *S_pd_* score, which we called the reference PD scores (*R_pd_*). For *R_pd_* we sampled 25 random reference SNP (SNPs that were not significant for any of the two GWA methods) and obtained a *R_pd_* distribution based on 1,000 *R_pd_* values. Following the concept of *S_pd_*, we also estimated a Kmer score (*K_pd_*), which consisted of the proportion of Kmers associated with resistance across populations. A value of 1 indicates that the individual has all the resistant Kmers and none of the susceptible Kmers (see Above), while a value of 0 indicates that the individual has all the susceptible kmers and none of the resistant ones. Finally, we also estimated a CNV score (*C_pd_*) that corresponds to the number of adaptive copy variants in an individual, where a higher number of CN variants indicates that the individual is more resistant to PD. For the *S_pd_*, *K_pd_* and *C_pd_* scores, we estimated the fit between the scores and the bacterial load. Finally, for the genetic independent variables, we also analyzed the linear model between the assignments into genetic groups (K1 and K2) based on the admixture analyses and the concentration of bacterial load.

### Climatic modeling

Given the predictive power of BIO8 and its general association with locations where *Xylella fastidiosa* has been detected, we used BIO8 to model the future distribution of *Xylella fastidiosa*. Assuming that 10°C is a predictive threshold of where *Xylella fastidiosa* is more likely to be present or absent, we identified areas in the globe that currently have BIO8 <10°C in the present but are predicted to have BIO8 >10°C in the future. We interpret these areas could become susceptible to *Xylella fastidiosa* in the future. We also detected the conversion - i.e. regions across the globe that currently have BIO8 >10°C in the present but are predicted to have BIO8 <10°C in the future - which we interpret as regions becoming less apt to harbor *Xylella fastidiosa* in the future. To make these predictions, we downloaded the BIO8 data at a 2.5 minutes resolution from Worldclim 2 (Fick and Hijmans 2017) for the present and for 54 climatic models in the future to consider the uncertainty in future climate projections. These future climate models included five global circulation models (GFDL-ESM4, IPSL-CM6A-LR, MPI-ESM1-2-HR, MRI-ESM2-0, UKESM1-0-LL), three time periods (2041-2060; 2061-2080; 2081-2100) and 4 shared socioeconomic pathways (SSPs: 126, 245, 370, 585). To plot the areas where climate is expected to change across the 10°C threshold, for the present and the 54 future layers we set the raster layers to 1 and 0 if they were above or below 10°C, respectively. Next, we subtracted each future layer to the present layer. Areas with values of −1 indicate that a region in the future layer is expected to be <10°C but is >10°C in the present (0-1). Regions with values of 0 indicate that both the present and future layers have values >10°C (1-1). Regions with values of +1 indicate that the future layer is expected to be >10°C but the present layer is <10°C (1-0). We did this for the 54 layers, and we created a sum_raster object by adding the 54 raster layers. Positive values (1 to 60) indicate the number of layers for which the prediction is that the region will become warmer than the 10°C threshold. Negative values (−1 to −60) indicate the number of future layers for which the prediction is that the region will become colder than the 10°C threshold. Values of 0 indicate that for all layers, the area is >10°C and will remain >10°C or that it is <10°C and will remain <10°C for all the layers.

Finally, we tested how BIO8 will change in the future in areas where *V. arizonica, V. vinifera* (grapevines), *Citrus* sp *Olea europea* (olives), *Prunnus amygdalus* (almond) and *Coffea* sp grow. For this, we first used the gbif function in the R dismo package (Hijmans et al.) to download all the known locations of V *arizonica*, grapevines, olives, almonds, coffee sp. and *Citrus* sp (gbif.org; download data: 2022-06-06). Next, for each species we used the CoordinateCleaner package in R (Zizka et al. 2019) to remove locations that 1) were duplicated; 2) had equal longitude and latitude; 3) were next to country centroids, capitals of countries, biological stations or gbif headquarters; 4) were in the sea; 5) were outliers based on the “quantile” option. Finally, we also removed locations if they were the only report in a given country, suggesting they may be outliers. After cleaning the data, we retained 6,204 locations (from 11,834) for *Coffee* sps.; 3,386 locations (from 9,992) for almonds; 1,111 locations (from 1,155) for *V. arizonica;* 5,256 locations (from 7,853) for *Citrus* sps.; 174,713 locations (from 204,775) for olives; and 33,225 locations (from 47,075) for grapevines.

For each species, we used the extract function in the raster R package (Hijmans 2021) to obtain for each location whether climate is expected to change across the 10°C layer (values −1,0 and 1; as detailed above). For each species we estimated the percentage of locations that are expected to move below and above the 10°C threshold across the 54 layers.

## ACKNOWLEDGEMENTS

The authors are grateful for data generation by R. Gaut and the Genomics High Throughput Facility at UC Irvine. D. Seymour and C. Roper at UC Riverside provided valuable comments. This work was funded by National Science Foundation US grant #1741627 to BSG, AW and DC.

