## Supplemental Tables and Figures for "Multigenic resistance to *Xylella fastidiosa* in wild grapes (*Vitis* sps.) and its implications within a changing climate"

**This PDF includes:**

Figs. S1 to S16
Tables S1 to S4
Tables S5 to S9 are available in separate *.xls files

SUPPLEMENTAL FIGURES

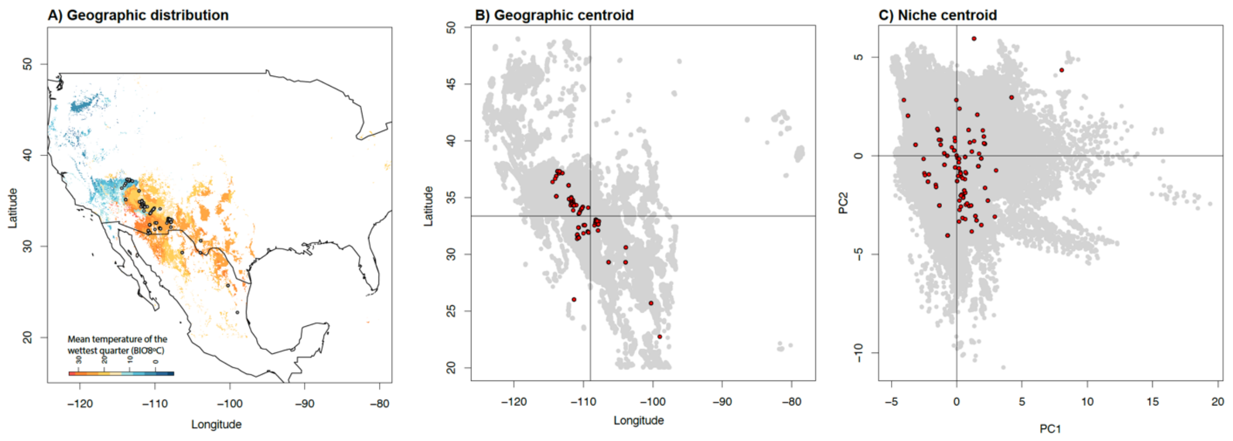


**Fig. S1.**

Geographic locations of *V. arizonica* samples used in this study (dots in each graph) illustrate a reasonable compared to A) the Species Distribution Model (SDM) projections for extant environments where the species could exist in theory based on BIO8, B) the predicted geographic centroid of the species and C) the predicted niche centroid of the species.

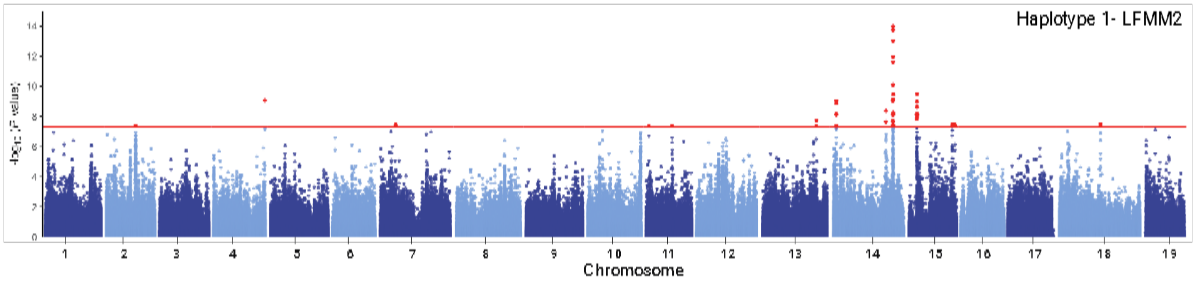


**Fig. S2.**
Manhattan plot from the genome-wide association studies of SNPs predicted in hap1 with *X. fastidiosa* loads (CFU/ml) performed by LFMM2. The red line indicates Bonferroni adjusted p-value of 0.05, which was used as the significance threshold.

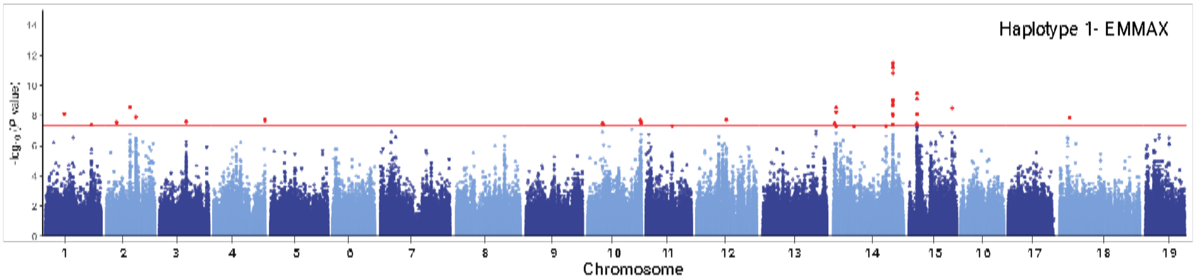


**Fig. S3.**

Manhattan plot from the genome-wide association studies of SNPs predicted in hap1 with *X. fastidiosa* loads (CFU/ml) performed by EMMAX. The red line indicates Bonferroni adjusted p-value of 0.05, which was used as the significance threshold.

A)

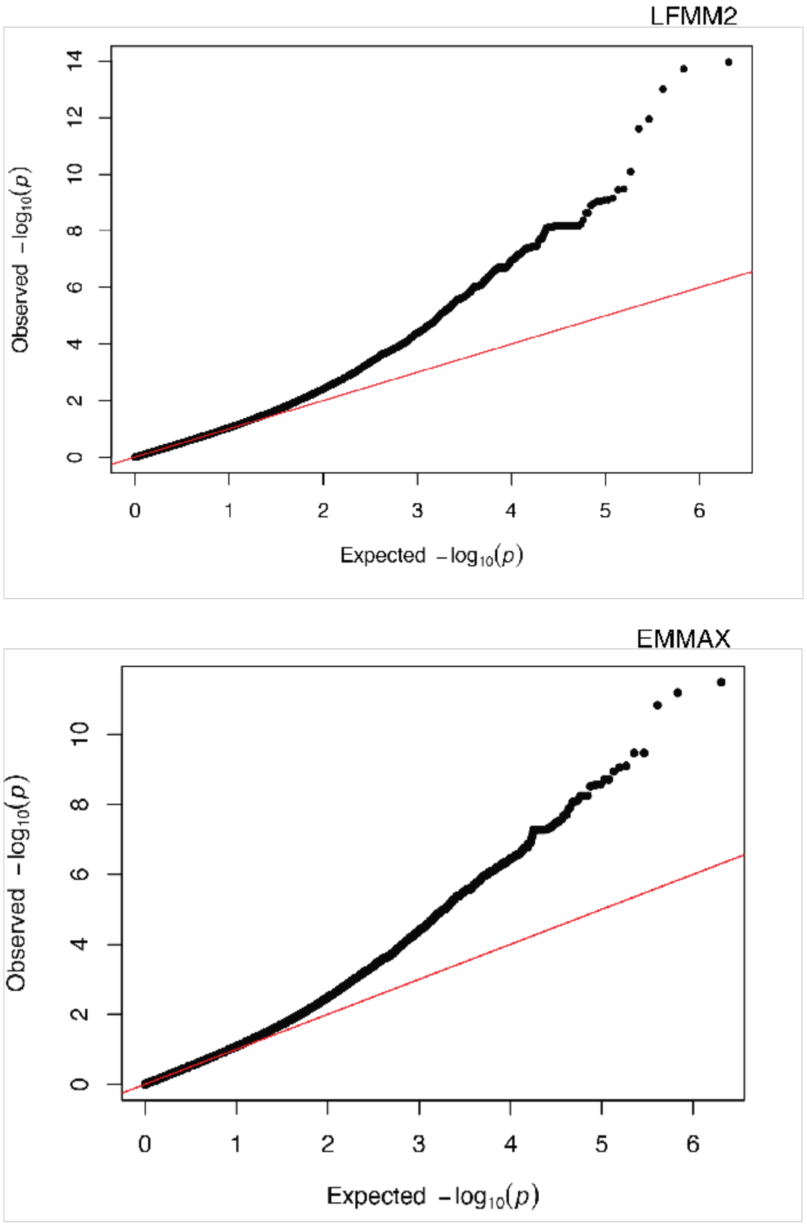


B)



**Fig. S4.**
Q-Q plot of p-values for the GWAS performed by A) LFMM2 and B) EMMAX on hap1.

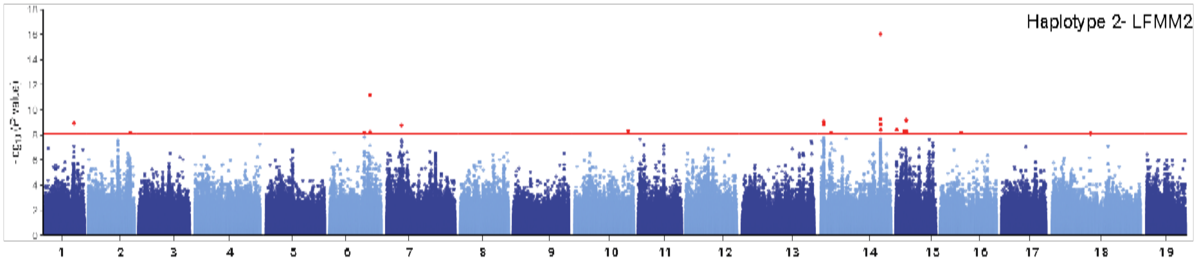


**Fig. S5.**

Manhattan plot from the genome-wide association studies of SNPs predicted in hap2 and *X. fastidiosa* loads (CFU/ml), as performed by LFMM2. The red line indicates Bonferroni adjusted p-value of 0.05, which was used as the significance threshold.

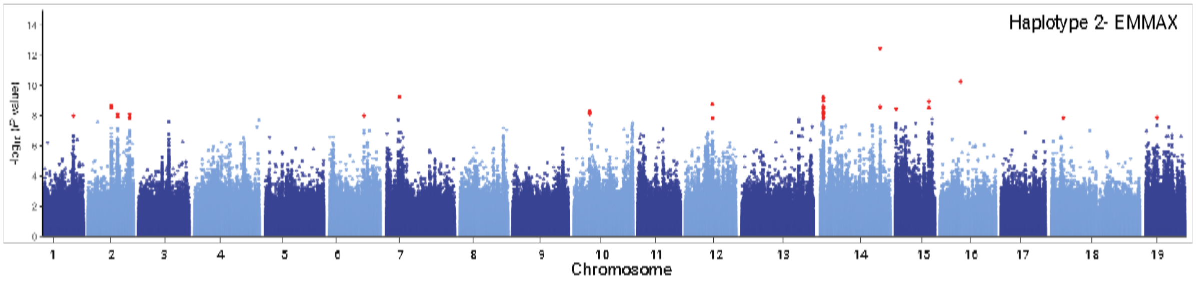


**Fig. S6.**

Manhattan plot from the genome-wide association studies of SNPs predicted in hap1 and *X. fastidiosa* loads (CFU/ml) performed by EMMAX. The red line indicates Bonferroni adjusted p-value of 0.05, which was used as the significance threshold.

A)

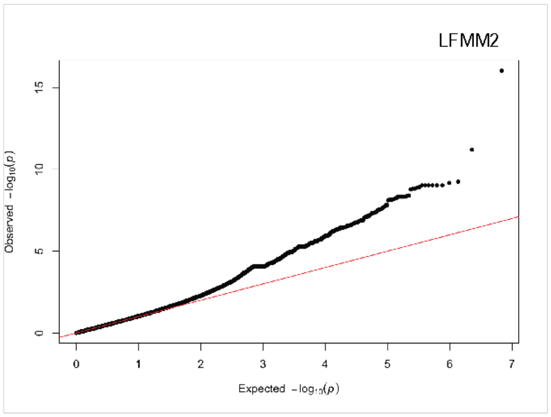


B)

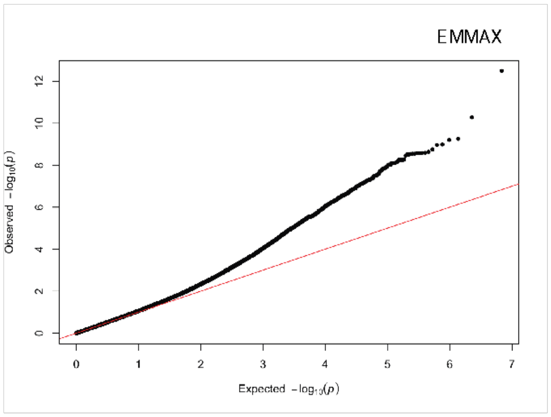


**Fig. S7.**
Q-Q plot of p-values for the GWAS in hap2 performed by A) LFMM2 and B) EMMAX.

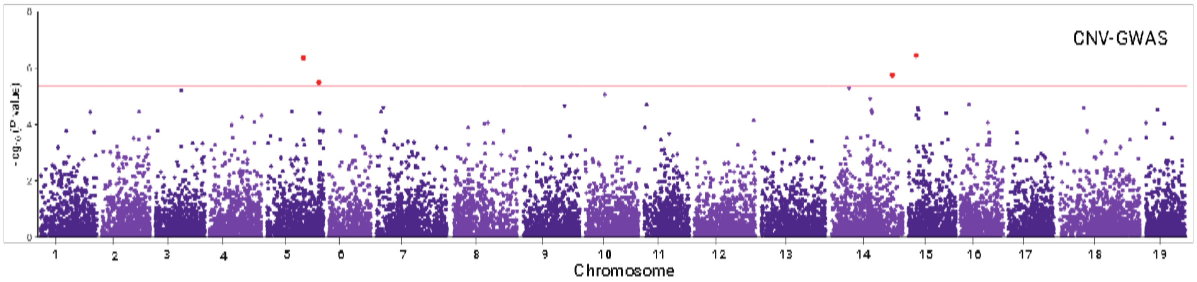


**Fig. S8.**

Manhattan plot from the genome-wide association studies of Copy Number Variants (CNVs) predicted in hap1 and *X. fastidiosa* loads (CFU/ml). The red line indicates Bonferroni adjusted p-value of 0.05, which was used as the significance threshold.

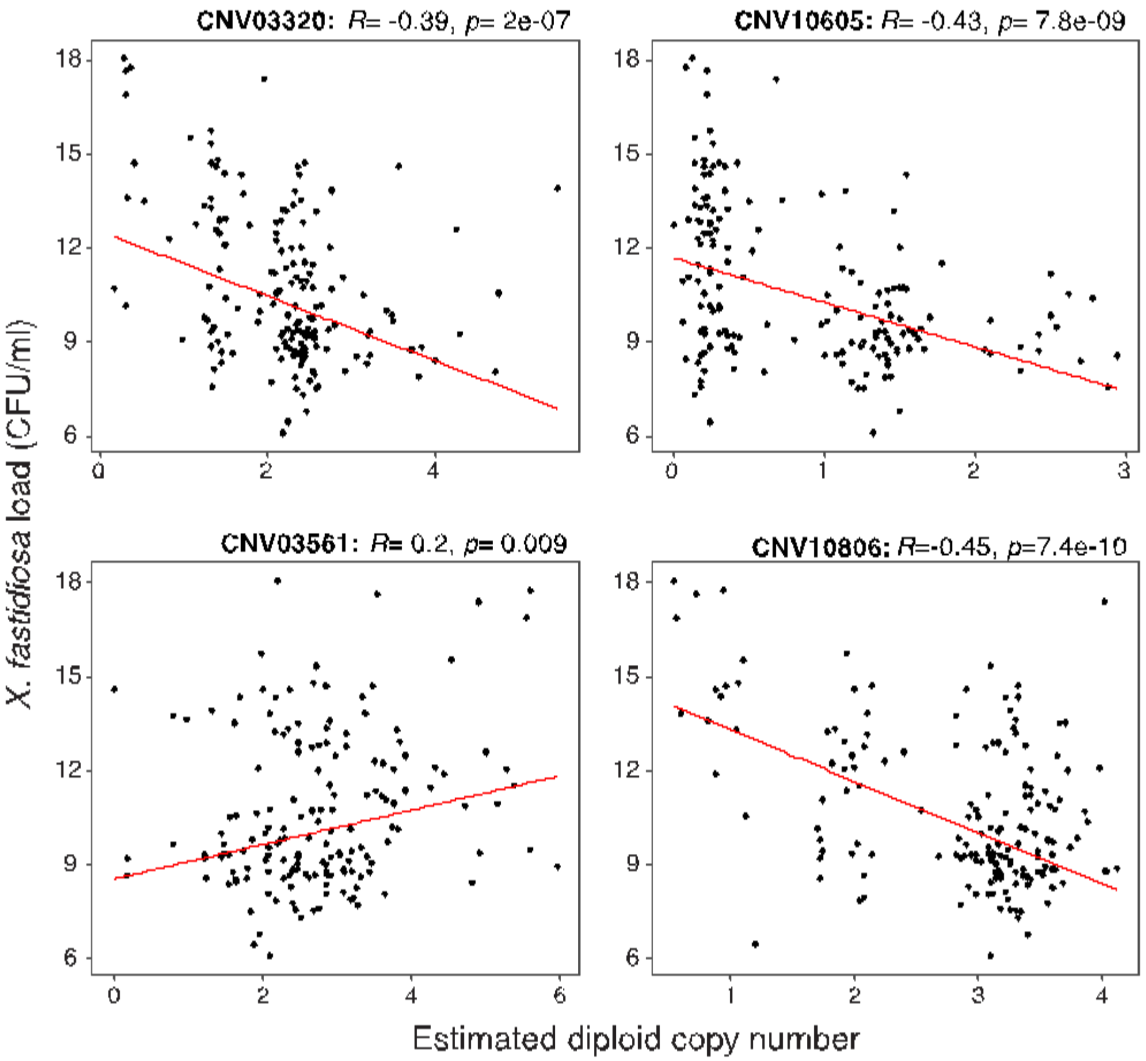


**Fig. S9.**

Scatterplots of four CNVs significantly associated with *X. fastidiosa* loads. Each dot represents an individual, with its estimated load (y-axis) and estimated diploid copy number (x-axis). Negative slopes indicate that higher copy numbers are associated with higher resistance.

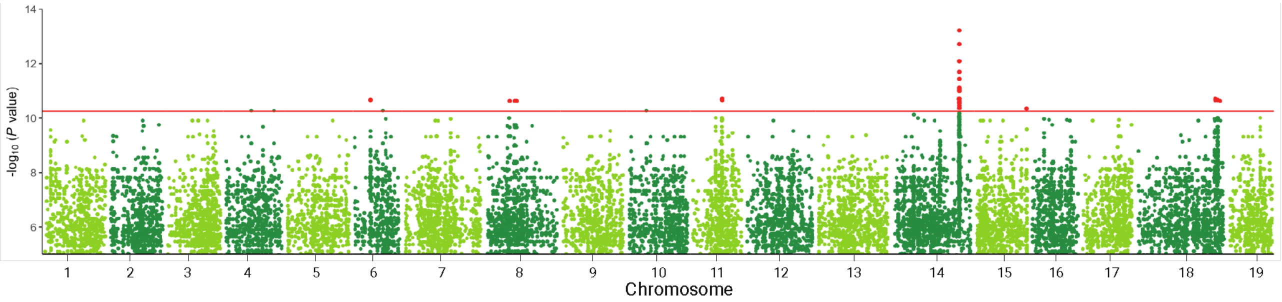


**Fig. S10.**

Manhattan plot from the genome-wide association studies of kmers with *X. fastidiosa* loads (CFU/ml). The plot shows kmers mapping to hap1 with a maximum of one mismatch. The red line indicates Bonferonni adjusted p-value of 0.05, which was used as the significance threshold.

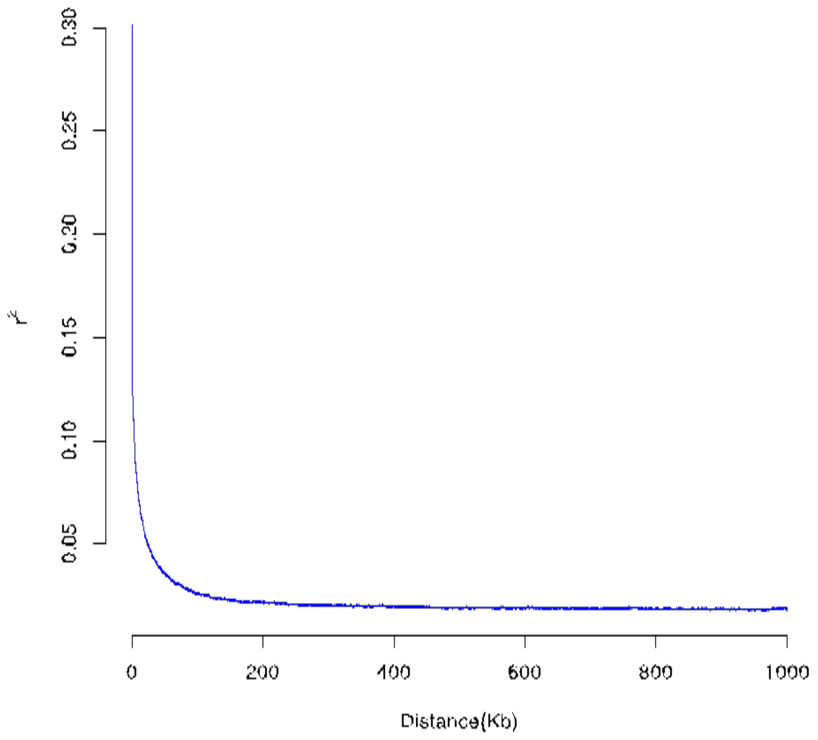


**Fig. S11.**
Estimated **l**inkage disequilibrium decay of genome-wide SNPs from the 167 *V. arizonica* accessions.

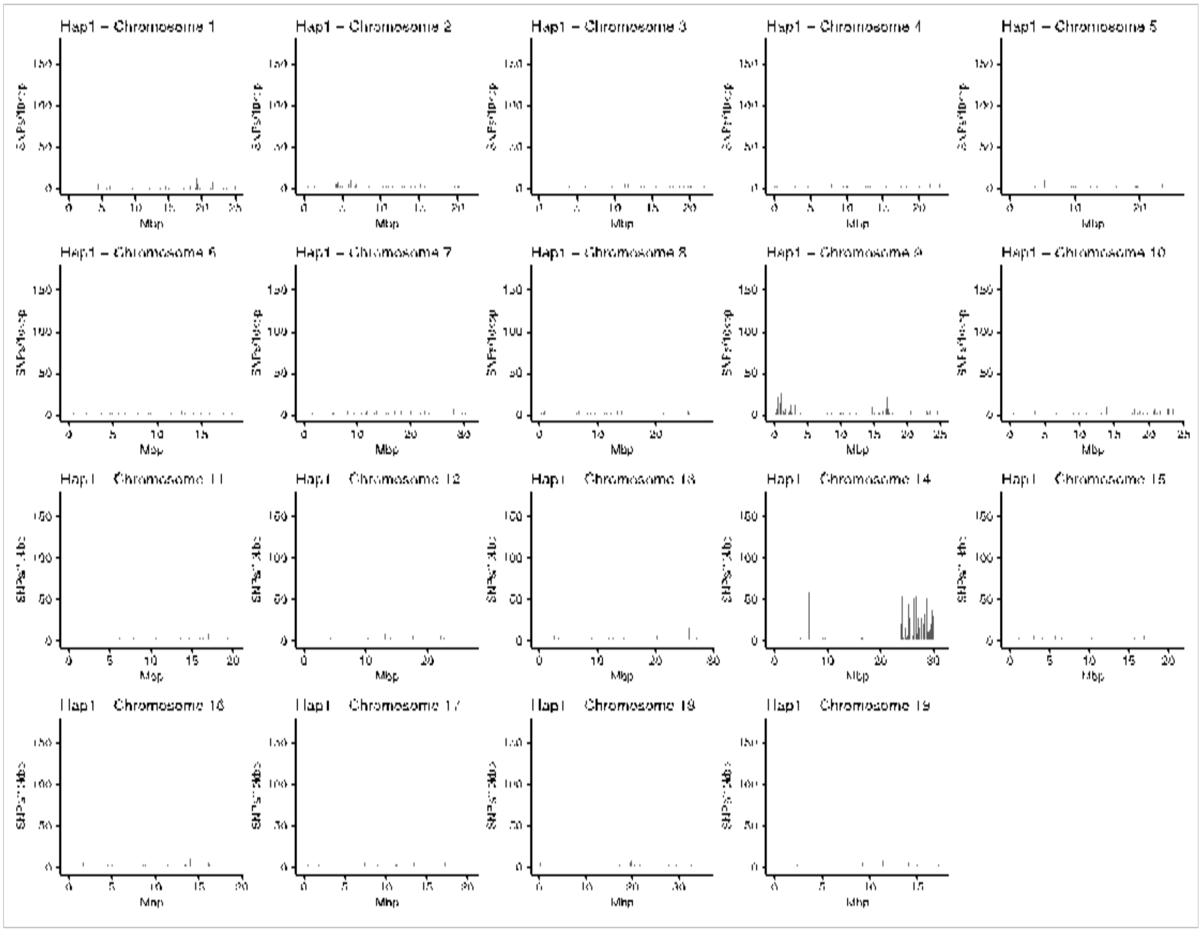


**Fig. S12.**
Plots of the 19 chromosomes, based on heterozygosity across 5 varietals backcrossed for PD resistance. The peaks represent the number of heterozygous SNPs across 10kb windows in which one of the heterozygous alleles was contributed by *V. arizonica*.

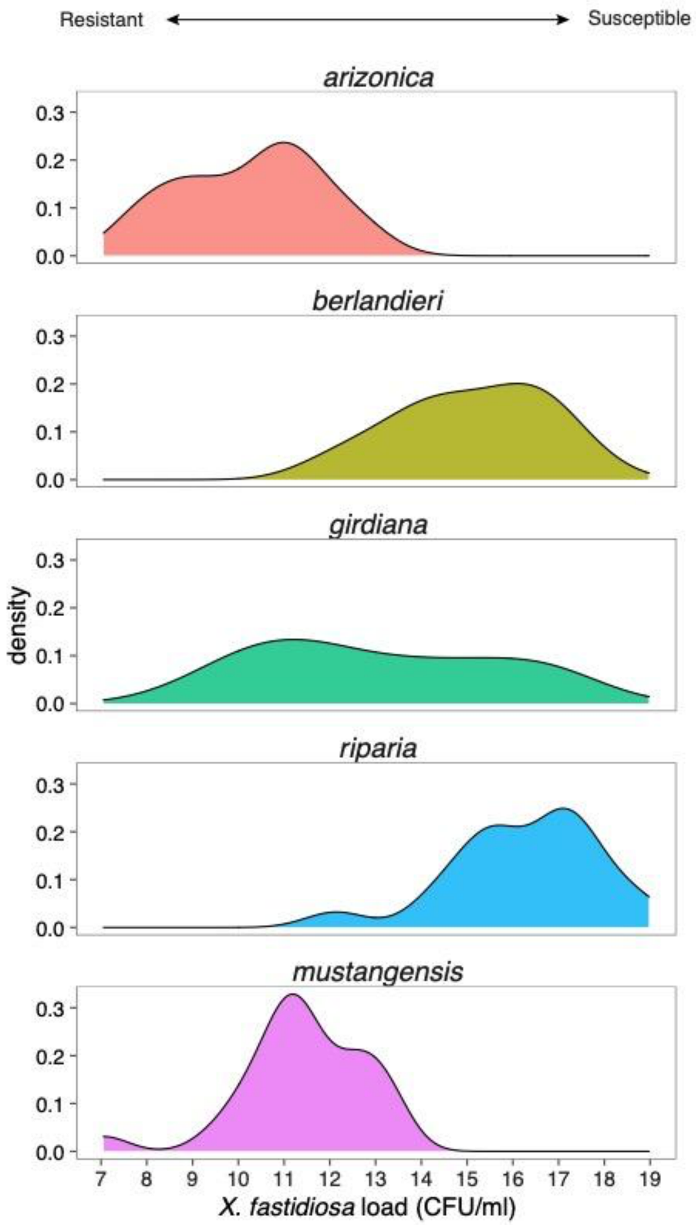


**Fig. S13.**
Distribution of *X. fastidiosa* loads (CFU/ml) in evaluated individuals of six wild grape species. Sample sizes for each species are provided in Fig. S14.

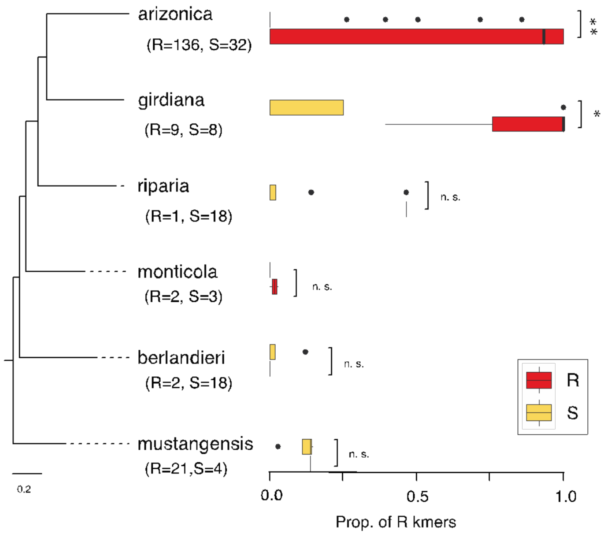


**Fig. S14.**

The frequencies of R-kmers in resistant (in red) and susceptible (yellow) individuals across six wild grape species. The sample sizes for resistant (R) and susceptible (S) individuals in each sample is provided in the paranthesis under species name. The inferred phylogeny of the species is provided to the left. Both *V. arizonica* and *V. girdiana* have significant differences in the frequency of R-kmers between R and S individuals, as indicated by asterisks on the right, but the other species do not have significant differences for R-kmers between R and S individuals. In fact, R-kmers are found less often than expected in *V. riparia*, *V. monticola*, *V. berlandieri* and *V. mustangensis* than control kmers chosen to have similar population frequencies in *V. arizonica* as R-kmers (Fig. 4C).

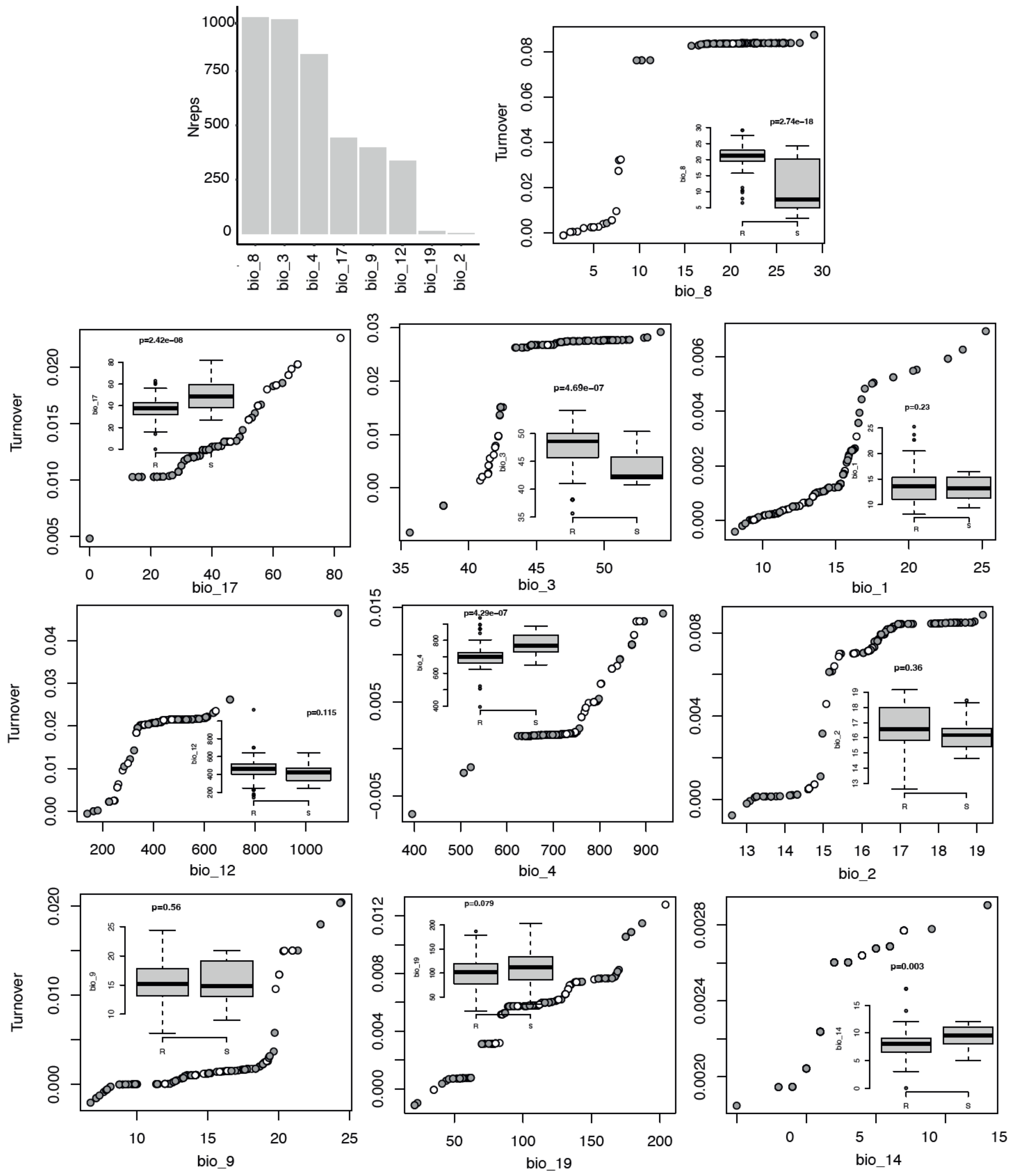


**Fig. S15.**
Results from Gradient Forest analyses. The top left graph is a histogram of the number of times, out of 1000 GF runs, that a particular bioclimatic variable was found to be among the top-3 most important variables. The remaining graphs show the turnover function for each of the 10 bioclimatic variables included in GF analyses, based on a single GF run. For each turnover graph, the x-axis is the distribution of the variable across sample sites (e.g., in degrees C for bio3 and several other variables.)

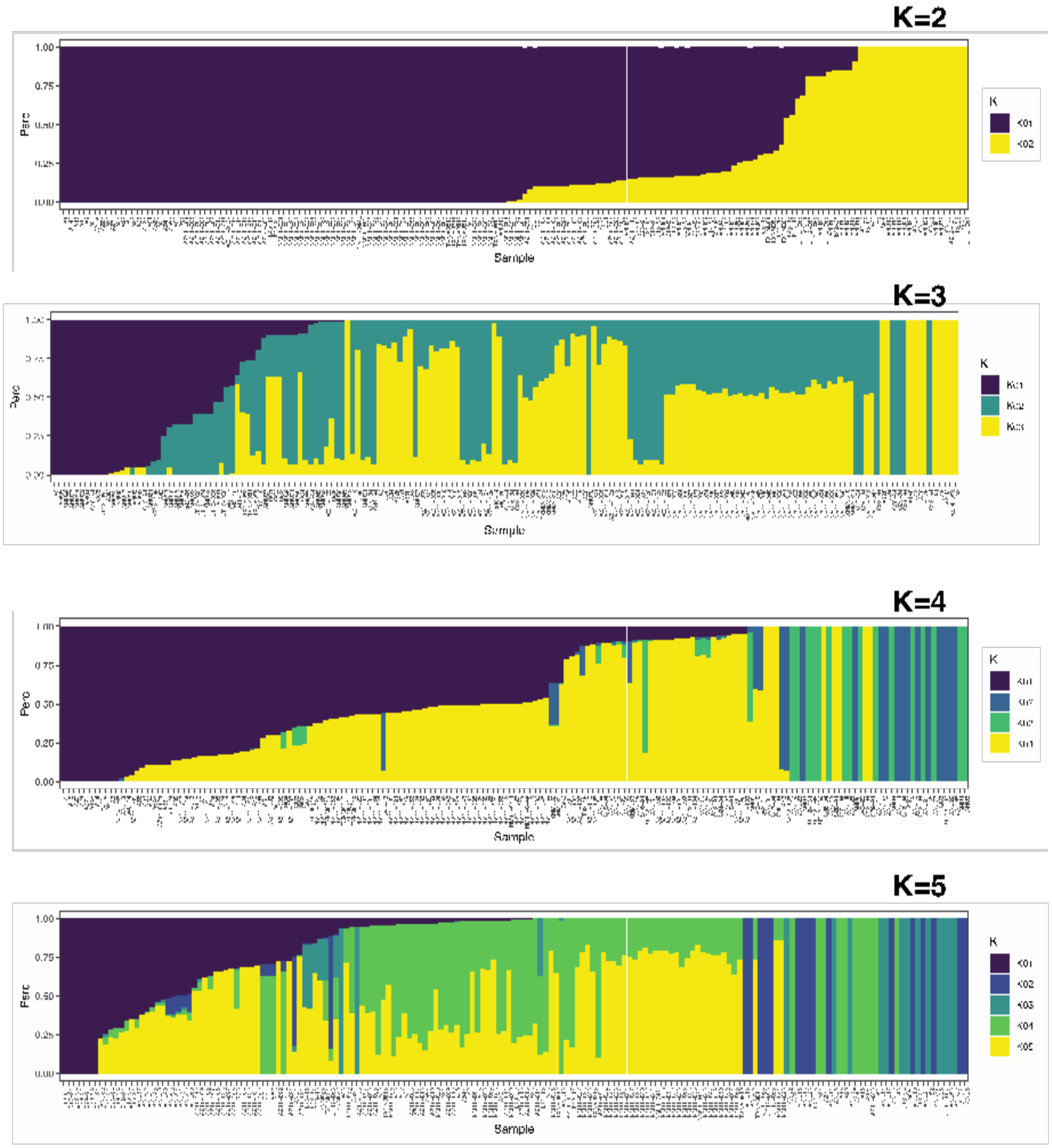


**Fig. S16.**

Genetic structure of *V. arizonica* with grouping values (*K*) of 2 to 5. The best grouping corresponds to K=2.

### SUPPLEMENTAL TABLES

**Table S1.**

Quantitative concentrations, in average CFU/ml across at least six replicates, of *X. fastidiosa* detected in the V. *arizonica* accessions used in this study. The data were originally reported in Morales-Cruz et al. (2021) and Riaz et al. (2020).

| No. | Sample | CFUs/ml |
|---|---|---|
| 1 | A1 | 8.691832 |
| 2 | A14F | 7.306167 |
| 3 | A15 | 9.231725 |
| 4 | A18 | 8.697725 |
| 5 | A2 | 7.758652 |
| 6 | A20 | 9.3398 |
| 7 | A21M | 9.095725 |
| 8 | A22F | 8.445193 |
| 9 | A24 | 9.139725 |
| 10 | A27F | 9.331725 |
| 11 | A28 | 9.786691 |
| 12 | A29 | 7.526725 |
| 13 | A3 | 8.761725 |
| 14 | A34 | 10.15700 |
| 15 | A35 | 10.68372 |
| 16 | A40 | 12.02772 |
| 17 | A42F | 6.774103 |
| 18 | A43 | 10.50972 |
| 19 | A5 | 9.214225 |
| 20 | A51 | 9.731003 |
| 21 | A52F | 8.645725 |
| 22 | A53 | 8.874248 |
| 23 | A54F | 9.323725 |
| 24 | A55F | 8.551725 |
| 25 | A56 | 8.067725 |
| 26 | A58 | 7.555725 |
| 27 | A8M | 8.050103 |
| 28 | ANU1 | 12.91549 |
| 29 | ANU10 | 17.64216 |
| 30 | ANU11 | 18.04422 |
| 31 | ANU13 | 14.32410 |
| 32 | ANU14 | 13.61216 |

| | | |
|---|---|---|
| 33 | ANU15 | 13.32590 |
| 34 | ANU16 | 14.69900 |
| 35 | ANU17 | 14.33210 |
| 36 | ANU18 | 15.72216 |
| 37 | ANU21 | 17.74896 |
| 38 | ANU23 | 14.35616 |
| 39 | ANU28 | 14.59549 |
| 40 | ANU29 | 13.90610 |
| 41 | ANU32 | 13.35972 |
| 42 | ANU4 | 10.51900 |
| 43 | ANU45 | 9.977725 |
| 44 | ANU49 | 11.21172 |
| 45 | ANU5 | 8.765345 |
| 46 | ANU50 | 13.82016 |
| 47 | ANU51 | 12.73216 |
| 48 | ANU52 | 13.51610 |
| 49 | ANU53 | 11.33298 |
| 50 | ANU57 | 9.417128 |
| 51 | ANU6 | 17.37852 |
| 52 | ANU65 | 12.10500 |
| 53 | ANU66 | 12.28839 |
| 54 | ANU67 | 10.14816 |
| 55 | ANU68 | 9.636103 |
| 56 | ANU69 | 10.71390 |
| 57 | ANU71 | 8.638167 |
| 58 | ANU77 | 8.754351 |
| 59 | ANU9 | 16.87216 |
| 60 | AZ11-001 | 9.149725 |
| 61 | AZ11-005 | 9.255058 |
| 62 | AZ11-006 | 9.291725 |
| 63 | AZ11-007 | 9.271725 |
| 64 | AZ11-008 | 9.299725 |
| 65 | AZ11-009 | 9.681906 |
| 66 | AZ11-010 | 9.195725 |
| 67 | AZ11-011 | 8.408103 |
| 68 | AZ11-012a | 8.939918 |
| 69 | AZ11-014 | 10.74900 |
| 70 | AZ11-016 | 11.99191 |
| 71 | AZ11-017 | 10.13372 |
| 72 | AZ11-018 | 10.12810 |

| | | |
|---|---|---|
| 73 | AZ11-019 | 8.559918 |
| 74 | AZ11-091 | 11.88772 |
| 75 | AZ11-092 | 12.46572 |
| 76 | AZ11-094 | 14.71258 |
| 77 | AZ11-096 | 15.33100 |
| 78 | AZ11-097 | 12.88791 |
| 79 | AZ11-099 | 15.51360 |
| 80 | AZ11-100 | 11.06172 |
| 81 | AZ11-101 | 13.80972 |
| 82 | AZ11-103 | 12.57772 |
| 83 | AZ11-104 | 10.04772 |
| 84 | AZ11-105 | 7.864225 |
| 85 | AZ11-107 | 8.297725 |
| 86 | AZ11-108 | 14.59700 |
| 87 | AZ11-109 | 9.061918 |
| 88 | AZ11-110 | 13.28777 |
| 89 | AZ12-138 | 14.59924 |
| 90 | AZ14-088 | 6.092103 |
| 91 | b40-14 | 9.844489 |
| 92 | b40-29 | 9.009259 |
| 93 | b41-13 | 7.514489 |
| 94 | b42-26 | 9.325414 |
| 95 | b43-15 | 8.387961 |
| 96 | b43-17 | 9.233416 |
| 97 | C12-94 | 8.119193 |
| 98 | C16-94 | 12.87008 |
| 99 | C17-94 | 6.446103 |
| 100 | C18-94 | 11.06024 |
| 101 | C19-94 | 11.49610 |
| 102 | C23-94 | 9.356167 |
| 103 | C24-94 | 8.849437 |
| 104 | C28-94 | 7.589193 |
| 105 | C29-94 | 8.346103 |
| 106 | C30-94 | 10.54519 |
| 107 | C31-94 | 9.297193 |
| 108 | C38-94 | 8.060103 |
| 109 | C42-94 | 12.46590 |
| 110 | GC1 | 8.535725 |
| 111 | GC2 | 9.987003 |
| 112 | GC3 | 8.567725 |

| | | |
|---|---|---|
| 113 | GC4 | 9.639003 |
| 114 | GC5 | 8.646103 |
| 115 | GC6 | 9.388442 |
| 116 | NM11-020 | 9.849003 |
| 117 | NM11-021 | 8.784918 |
| 118 | NM11-022 | 12.79810 |
| 119 | NM11-023 | 10.63422 |
| 120 | NM11-024 | 11.22922 |
| 121 | NM11-026 | 8.773918 |
| 122 | NM11-027 | 10.75991 |
| 123 | NM11-031 | 7.942103 |
| 124 | NM11-033 | 10.74191 |
| 125 | NM11-034 | 12.76210 |
| 126 | NM11-035 | 10.93140 |
| 127 | NM11-036 | 7.707725 |
| 128 | NM11-037 | 10.21100 |
| 129 | NM11-038 | 8.845087 |
| 130 | NM11-039 | 12.22772 |
| 131 | NM11-040 | 11.45791 |
| 132 | NM11-042 | 11.34991 |
| 133 | NM11-043 | 9.465918 |
| 134 | NM11-044a | 9.235918 |
| 135 | NM11-044f | 7.889725 |
| 136 | NM11-045 | 11.18172 |
| 137 | NM11-046 | 10.87100 |
| 138 | NM11-047 | 11.53491 |
| 139 | NM11-048 | 8.603603 |
| 140 | NM11-049 | 9.553725 |
| 141 | NM11-050 | 8.667418 |
| 142 | NM11-051 | 10.38591 |
| 143 | NM11-052 | 9.657003 |
| 144 | NM11-053 | 13.14860 |
| 145 | NM11-055 | 12.03191 |
| 146 | NM11-058 | 8.801918 |
| 147 | NM11-059 | 9.772587 |
| 148 | NM11-060 | 12.58010 |
| 149 | NM11-061 | 12.80972 |
| 150 | NM11-063 | 13.18325 |
| 151 | NM11-064 | 9.423725 |
| 152 | NM11-065 | 8.981725 |

| | | |
|---|---|---|
| 153 | NM11-066 | 8.839725 |
| 154 | NM11-067 | 9.546725 |
| 155 | NM11-068 | 9.104918 |
| 156 | TXNM081 | 13.49216 |
| 157 | TXNM0810 | 8.593003 |
| 158 | TXNM0811 | 8.258527 |
| 159 | TXNM0812 | 12.07313 |
| 160 | TXNM0813 | 8.768392 |
| 161 | TXNM0816 | 10.93610 |
| 162 | TXNM082 | 14.69772 |
| 163 | TXNM086 | 10.54590 |
| 164 | UT12-065 | 14.79100 |
| 165 | UT12-066 | 13.57140 |
| 166 | UT12-067 | 13.72124 |
| 167 | UT12-068 | 13.22900 |

**Table S2.**
Significant SNPs from the GWAS in hap1 and hap2 and from both methods.

| Haplotype | SNP_ID | chr | pos | LFMM (pval) | EMMAX (pval) |
|---|---|---|---|---|---|
| hap1 | chr01_8370148 | chr01 | 8370148 | N.S. | 7.38E-09 |
| hap1 | chr01_20596920 | chr01 | 20596920 | N.S. | 4.11E-08 |
| hap1 | chr02_4485424 | chr02 | 4485424 | N.S. | 2.72E-08 |
| hap1 | chr02_10644429 | chr02 | 10644429 | N.S. | 2.69E-09 |
| hap1 | chr02_10644444 | chr02 | 10644444 | N.S. | 2.69E-09 |
| hap1 | chr02_13077962 | chr02 | 13077962 | 4.09E-08 | N.S. |
| hap1 | chr02_13108108 | chr02 | 13108108 | 4.09E-08 | N.S. |
| hap1 | chr02_13149072 | chr02 | 13149072 | 4.09E-08 | 1.22E-08 |
| hap1 | chr02_13149103 | chr02 | 13149103 | 4.09E-08 | N.S. |
| hap1 | chr03_11989991 | chr03 | 11989991 | N.S. | 2.51E-08 |
| hap1 | chr04_22919135 | chr04 | 22919135 | 8.29E-10 | 1.98E-08 |
| hap1 | chr07_6871078 | chr07 | 6871078 | 3.53E-08 | N.S. |
| hap1 | chr07_6871526 | chr07 | 6871526 | 3.53E-08 | N.S. |
| hap1 | chr10_6500495 | chr10 | 6500495 | N.S. | 3.19E-08 |
| hap1 | chr10_7085360 | chr10 | 7085360 | N.S. | 4.00E-08 |
| hap1 | chr10_23462586 | chr10 | 23462586 | N.S. | 2.05E-08 |
| hap1 | chr10_23765386 | chr10 | 23765386 | N.S. | 2.88E-08 |
| hap1 | chr11_1230838 | chr11 | 1230838 | 4.26E-08 | N.S. |
| hap1 | chr11_11647678 | chr11 | 11647678 | 4.09E-08 | 4.54E-08 |
| hap1 | chr12_13334771 | chr12 | 13334771 | N.S. | 1.92E-08 |
| hap1 | chr13_23870570 | chr13 | 23870570 | 4.26E-08 | N.S. |
| hap1 | chr13_23915236 | chr13 | 23915236 | 1.83E-08 | N.S. |
| hap1 | chr14_475430 | chr14 | 475430 | N.S. | 3.29E-08 |
| hap1 | chr14_1234136 | chr14 | 1234136 | 7.54E-09 | 5.63E-09 |
| hap1 | chr14_1235411 | chr14 | 1235411 | 4.71E-08 | N.S. |
| hap1 | chr14_1235777 | chr14 | 1235777 | 9.02E-10 | 4.78E-08 |
| hap1 | chr14_1235915 | chr14 | 1235915 | 7.54E-09 | 5.63E-09 |
| hap1 | chr14_1236268 | chr14 | 1236268 | 1.27E-09 | 2.89E-09 |
| hap1 | chr14_1236972 | chr14 | 1236972 | 7.54E-09 | 5.63E-09 |
| hap1 | chr14_1237202 | chr14 | 1237202 | 7.54E-09 | 5.63E-09 |
| hap1 | chr14_1237928 | chr14 | 1237928 | 3.95E-08 | N.S. |
| hap1 | chr14_1237998 | chr14 | 1237998 | 6.72E-09 | N.S. |
| hap1 | chr14_9132810 | chr14 | 9132810 | N.S. | 4.90E-08 |
| hap1 | chr14_23480870 | chr14 | 23480870 | 2.19E-08 | N.S. |

| | | | | | |
|---|---|---|---|---|---|
| hap1 | chr14_23506264 | chr14 | 23506264 | 4.13E-09 | 4.54E-08 |
| hap1 | chr14_26593134 | chr14 | 26593134 | 1.89E-08 | N.S. |
| hap1 | chr14_26593913 | chr14 | 26593913 | 1.89E-08 | N.S. |
| hap1 | chr14_26598435 | chr14 | 26598435 | 9.05E-10 | 8.25E-09 |
| hap1 | chr14_26632319 | chr14 | 26632319 | 7.92E-09 | 1.94E-09 |
| hap1 | chr14_26632651 | chr14 | 26632651 | N.S. | 3.72E-08 |
| hap1 | chr14_26637424 | chr14 | 26637424 | 1.89E-08 | N.S. |
| hap1 | chr14_26641323 | chr14 | 26641323 | 7.92E-09 | 1.94E-09 |
| hap1 | chr14_26650901 | chr14 | 26650901 | 7.97E-10 | N.S. |
| hap1 | chr14_26657700 | chr14 | 26657700 | 2.28E-08 | N.S. |
| hap1 | chr14_26657901 | chr14 | 26657901 | 3.72E-08 | N.S. |
| hap1 | chr14_26658196 | chr14 | 26658196 | 8.13E-11 | 9.12E-09 |
| hap1 | chr14_26674335 | chr14 | 26674335 | 3.66E-08 | N.S. |
| hap1 | chr14_26677939 | chr14 | 26677939 | 1.08E-14 | 3.21E-12 |
| hap1 | chr14_26678120 | chr14 | 26678120 | 2.46E-12 | 8.68E-10 |
| hap1 | chr14_26679420 | chr14 | 26679420 | 1.13E-12 | 1.13E-09 |
| hap1 | chr14_26680207 | chr14 | 26680207 | 1.88E-14 | 6.43E-12 |
| hap1 | chr14_26682193 | chr14 | 26682193 | 7.21E-09 | N.S. |
| hap1 | chr14_26684394 | chr14 | 26684394 | 7.02E-10 | N.S. |
| hap1 | chr14_26684487 | chr14 | 26684487 | 9.69E-14 | 1.46E-11 |
| hap1 | chr14_26701723 | chr14 | 26701723 | 4.14E-08 | N.S. |
| hap1 | chr14_26733459 | chr14 | 26733459 | 5.67E-09 | N.S. |
| hap1 | chr14_26745142 | chr14 | 26745142 | 3.60E-08 | N.S. |
| hap1 | chr14_26752710 | chr14 | 26752710 | 7.40E-09 | N.S. |
| hap1 | chr14_26753827 | chr14 | 26753827 | 3.52E-10 | N.S. |
| hap1 | chr14_26759896 | chr14 | 26759896 | 3.87E-08 | N.S. |
| hap1 | chr15_3306840 | chr15 | 3306840 | 1.41E-08 | 3.35E-08 |
| hap1 | chr15_3310907 | chr15 | 3310907 | 6.78E-09 | N.S. |
| hap1 | chr15_3311798 | chr15 | 3311798 | 6.78E-09 | N.S. |
| hap1 | chr15_3311954 | chr15 | 3311954 | 6.78E-09 | N.S. |
| hap1 | chr15_3311975 | chr15 | 3311975 | 6.78E-09 | N.S. |
| hap1 | chr15_3373418 | chr15 | 3373418 | 1.18E-08 | N.S. |
| hap1 | chr15_3375239 | chr15 | 3375239 | 6.78E-09 | N.S. |
| hap1 | chr15_3375879 | chr15 | 3375879 | 2.31E-09 | 3.39E-10 |
| hap1 | chr15_3376889 | chr15 | 3376889 | 6.78E-09 | N.S. |
| hap1 | chr15_3377201 | chr15 | 3377201 | 6.78E-09 | N.S. |
| hap1 | chr15_3377369 | chr15 | 3377369 | 6.78E-09 | N.S. |
| hap1 | chr15_3377605 | chr15 | 3377605 | 6.78E-09 | N.S. |
| hap1 | chr15_3378559 | chr15 | 3378559 | 1.04E-09 | 7.99E-10 |
| hap1 | chr15_3382242 | chr15 | 3382242 | 6.78E-09 | N.S. |

| | | | | | |
|---|---|---|---|---|---|
| hap1 | chr15_3382625 | chr15 | 3382625 | 6.78E-09 | N.S. |
| hap1 | chr15_3383126 | chr15 | 3383126 | 6.78E-09 | N.S. |
| hap1 | chr15_3388120 | chr15 | 3388120 | 6.78E-09 | N.S. |
| hap1 | chr15_3389507 | chr15 | 3389507 | 2.31E-09 | 3.39E-10 |
| hap1 | chr15_3406925 | chr15 | 3406925 | 3.32E-10 | 8.10E-09 |
| hap1 | chr15_3407357 | chr15 | 3407357 | 6.78E-09 | N.S. |
| hap1 | chr15_3412870 | chr15 | 3412870 | 1.04E-08 | N.S. |
| hap1 | chr15_3520061 | chr15 | 3520061 | 6.78E-09 | N.S. |
| hap1 | chr15_3520064 | chr15 | 3520064 | 6.78E-09 | N.S. |
| hap1 | chr15_19131878 | chr15 | 19131878 | 3.32E-08 | 3.05E-09 |
| hap1 | chr15_20295279 | chr15 | 20295279 | 3.53E-08 | N.S. |
| hap1 | chr15_20295781 | chr15 | 20295781 | 3.53E-08 | N.S. |
| hap1 | chr15_20298765 | chr15 | 20298765 | 3.53E-08 | N.S. |
| hap1 | chr18_4471198 | chr18 | 4471198 | N.S. | 1.35E-08 |
| hap1 | chr18_18290018 | chr18 | 18290018 | 3.14E-08 | N.S. |
| hap2 | chr01_11713395 | chr01 | 11713395 | 1.18799E-09 | 9.62078E-09 |
| hap2 | chr02_9433839 | chr02 | 9433839 | N.S. | 2.68547E-09 |
| hap2 | chr02_9433854 | chr02 | 9433854 | N.S. | 2.68547E-09 |
| hap2 | chr02_9434049 | chr02 | 9434049 | N.S. | 2.33624E-09 |
| hap2 | chr02_11996700 | chr02 | 11996700 | N.S. | 1.21789E-08 |
| hap2 | chr02_11996746 | chr02 | 11996746 | N.S. | 7.73851E-09 |
| hap2 | chr02_11996766 | chr02 | 11996766 | N.S. | 7.73851E-09 |
| hap2 | chr02_12004337 | chr02 | 12004337 | N.S. | 1.15926E-08 |
| hap2 | chr02_16990744 | chr02 | 16990744 | 6.6269E-09 | 7.75219E-09 |
| hap2 | chr02_16992826 | chr02 | 16992826 | N.S. | 1.43276E-08 |
| hap2 | chr06_14076729 | chr06 | 14076729 | 7.25557E-09 | N.S. |
| hap2 | chr06_14076782 | chr06 | 14076782 | 7.25557E-09 | N.S. |
| hap2 | chr06_14076799 | chr06 | 14076799 | 7.25557E-09 | N.S. |
| hap2 | chr06_14077985 | chr06 | 14077985 | N.S. | 9.50843E-09 |
| hap2 | chr06_14077988 | chr06 | 14077988 | N.S. | 9.50843E-09 |
| hap2 | chr06_14077991 | chr06 | 14077991 | N.S. | 9.50843E-09 |
| hap2 | chr06_16352123 | chr06 | 16352123 | 5.83667E-09 | N.S. |
| hap2 | chr06_16352128 | chr06 | 16352128 | 6.21838E-12 | N.S. |
| hap2 | chr06_16352159 | chr06 | 16352159 | 5.83667E-09 | N.S. |
| hap2 | chr07_5349315 | chr07 | 5349315 | N.S. | 5.55425E-10 |
| hap2 | chr07_5940428 | chr07 | 5940428 | 1.6557E-09 | N.S. |
| hap2 | chr10_6353390 | chr10 | 6353390 | N.S. | 7.52371E-09 |
| hap2 | chr10_6354278 | chr10 | 6354278 | N.S. | 5.24863E-09 |
| hap2 | chr10_21695581 | chr10 | 21695581 | 4.76515E-09 | N.S. |
| hap2 | chr12_10915403 | chr12 | 10915403 | N.S. | 1.75734E-09 |

| | | | | | |
|---|---|---|---|---|---|
| hap2 | chr12_11027195 | chr12 | 11027195 | N.S. | 1.44863E-08 |
| hap2 | chr14_1153842 | chr14 | 1153842 | N.S. | 7.95516E-09 |
| hap2 | chr14_1156046 | chr14 | 1156046 | N.S. | 6.24054E-10 |
| hap2 | chr14_1156099 | chr14 | 1156099 | N.S. | 1.01911E-09 |
| hap2 | chr14_1156131 | chr14 | 1156131 | N.S. | 2.5432E-09 |
| hap2 | chr14_1156136 | chr14 | 1156136 | N.S. | 6.65157E-09 |
| hap2 | chr14_1156400 | chr14 | 1156400 | N.S. | 2.83958E-09 |
| hap2 | chr14_1156415 | chr14 | 1156415 | N.S. | 2.83958E-09 |
| hap2 | chr14_1172804 | chr14 | 1172804 | 9.16853E-10 | 5.62546E-09 |
| hap2 | chr14_1174015 | chr14 | 1174015 | 9.16853E-10 | 1.36385E-08 |
| hap2 | chr14_1174584 | chr14 | 1174584 | 9.16853E-10 | 5.62546E-09 |
| hap2 | chr14_1174834 | chr14 | 1174834 | 1.53884E-09 | N.S. |
| hap2 | chr14_1174937 | chr14 | 1174937 | N.S. | 2.88641E-09 |
| hap2 | chr14_1174994 | chr14 | 1174994 | 9.16853E-10 | 7.15451E-09 |
| hap2 | chr14_1175641 | chr14 | 1175641 | 9.16853E-10 | 5.62546E-09 |
| hap2 | chr14_1175871 | chr14 | 1175871 | 9.16853E-10 | 5.62546E-09 |
| hap2 | chr14_4184761 | chr14 | 4184761 | 7.25557E-09 | N.S. |
| hap2 | chr14_24218653 | chr14 | 24218653 | 9.135E-17 | 3.20754E-13 |
| hap2 | chr14_24231510 | chr14 | 24231510 | N.S. | 2.59794E-09 |
| hap2 | chr14_24244969 | chr14 | 24244969 | 5.62837E-10 | N.S. |
| hap2 | chr14_24306913 | chr14 | 24306913 | 4.14109E-09 | N.S. |
| hap2 | chr14_24316271 | chr14 | 24316271 | 1.53235E-09 | N.S. |
| hap2 | chr14_24317392 | chr14 | 24317392 | 1.31207E-09 | N.S. |
| hap2 | chr15_251538 | chr15 | 251538 | 3.97002E-09 | 3.45546E-09 |
| hap2 | chr15_3268733 | chr15 | 3268733 | 4.76515E-09 | N.S. |
| hap2 | chr15_3945043 | chr15 | 3945043 | 4.76515E-09 | N.S. |
| hap2 | chr15_3956390 | chr15 | 3956390 | 4.76515E-09 | N.S. |
| hap2 | chr15_3967765 | chr15 | 3967765 | 4.76515E-09 | N.S. |
| hap2 | chr15_3967773 | chr15 | 3967773 | 4.76515E-09 | N.S. |
| hap2 | chr15_4059655 | chr15 | 4059655 | 6.68405E-10 | N.S. |
| hap2 | chr15_4064825 | chr15 | 4064825 | 4.76515E-09 | N.S. |
| hap2 | chr15_4068329 | chr15 | 4068329 | 6.58886E-09 | N.S. |
| hap2 | chr15_13544980 | chr15 | 13544980 | N.S. | 3.05263E-09 |
| hap2 | chr15_13549576 | chr15 | 13549576 | N.S. | 1.07472E-09 |
| hap2 | chr15_13595853 | chr15 | 13595853 | N.S. | 3.11867E-09 |
| hap2 | chr16_8406896 | chr16 | 8406896 | 6.47001E-09 | 5.28713E-11 |
| hap2 | chr18_4727166 | chr18 | 4727166 | N.S. | 1.34935E-08 |
| hap2 | chr18_15550621 | chr18 | 15550621 | 7.78935E-09 | N.S. |
| hap2 | chr19_4472898 | chr19 | 4472898 | N.S. | 1.32508E-08 |

**Table S3.**
Copy number variants (CNVs) within the SNP-defined peaks associated with PD resistance.

| Peak | Chromosome | Copy Number Variant ID | Start | End | Size (bp) |
|---|---|---|---|---|---|
| 1 | VITVarB40-14_v2.0.hap1.chr02 | CNV01053 | 13094001 | 13098000 | 4000 |
| 1 | VITVarB40-14_v2.0.hap1.chr02 | CNV01054 | 13099001 | 13107000 | 8000 |
| 1 | VITVarB40-14_v2.0.hap1.chr02 | CNV01055 | 13108001 | 13115000 | 7000 |
| 1 | VITVarB40-14_v2.0.hap1.chr02 | CNV01051 | 13069001 | 13083000 | 14000 |
| 1 | VITVarB40-14_v2.0.hap1.chr02 | CNV01052 | 13084001 | 13093000 | 9000 |
| 1 | VITVarB40-14_v2.0.hap1.chr02 | CNV01056 | 13117001 | 13125000 | 8000 |
| 1 | VITVarB40-14_v2.0.hap1.chr02 | CNV01057 | 13128001 | 13150000 | 22000 |
| 1 | VITVarB40-14_v2.0.hap1.chr02 | CNV01058 | 13170001 | 13175000 | 5000 |
| 1 | VITVarB40-14_v2.0.hap1.chr02 | CNV01050 | 12955001 | 13057000 | 7929 |
| 1 | VITVarB40-14_v2.0.hap1.chr02 | CNV01059 | 13190001 | 13246000 | 56000 |
| 2 | VITVarB40-14_v2.0.hap1.chr04 | CNV02847 | 22938001 | 22945000 | 7000 |
| 2 | VITVarB40-14_v2.0.hap1.chr04 | CNV02848 | 22947001 | 22952000 | 5000 |
| 2 | VITVarB40-14_v2.0.hap1.chr04 | CNV02846 | 22866001 | 22876000 | 10000 |
| 2 | VITVarB40-14_v2.0.hap1.chr04 | CNV02849 | 22952001 | 22957000 | 5000 |
| 2 | VITVarB40-14_v2.0.hap1.chr04 | CNV02850 | 22962001 | 22974000 | 12000 |
| 2 | VITVarB40-14_v2.0.hap1.chr04 | CNV02851 | 22976001 | 23002000 | 26000 |
| 3 | VITVarB40-14_v2.0.hap1.chr11 | CNV07678 | 11552001 | 11564000 | 12000 |
| 3 | VITVarB40-14_v2.0.hap1.chr11 | CNV07683 | 11688001 | 11692000 | 4000 |
| 3 | VITVarB40-14_v2.0.hap1.chr11 | CNV07684 | 11723001 | 11727000 | 4000 |

| | | | | | |
|---|---|---|---|---|---|
| 3 | VITVarB40-14_v2.0.hap1.chr11 | CNV07685 | 11731001 | 11746000 | 15000 |
| 3 | VITVarB40-14_v2.0.hap1.chr11 | CNV07679 | 11563001 | 11577000 | 14000 |
| 3 | VITVarB40-14_v2.0.hap1.chr11 | CNV07680 | 11587001 | 11655000 | 68000 |
| 3 | VITVarB40-14_v2.0.hap1.chr11 | CNV07682 | 11680001 | 11686000 | 6000 |
| 3 | VITVarB40-14_v2.0.hap1.chr11 | CNV07681 | 11655001 | 11676000 | 21000 |
| 4 | VITVarB40-14_v2.0.hap1.chr14 | CNV09761 | 1186001 | 1196000 | 10000 |
| 4 | VITVarB40-14_v2.0.hap1.chr14 | CNV09762 | 1229001 | 1234000 | 5000 |
| 4 | VITVarB40-14_v2.0.hap1.chr14 | CNV09763 | 1252001 | 1261000 | 9000 |
| 5 | VITVarB40-14_v2.0.hap1.chr14 | CNV10539 | 23406001 | 23411000 | 4737 |
| 5 | VITVarB40-14_v2.0.hap1.chr14 | CNV10540 | 23437001 | 23443000 | 6000 |
| 5 | VITVarB40-14_v2.0.hap1.chr14 | CNV10541 | 23518001 | 23525000 | 7000 |
| 6 | VITVarB40-14_v2.0.hap1.chr14 | CNV10599 | 26611001 | 26616000 | 5000 |
| 6 | VITVarB40-14_v2.0.hap1.chr14 | CNV10600 | 26615001 | 26622000 | 7000 |
| 6 | VITVarB40-14_v2.0.hap1.chr14 | CNV10601 | 26626001 | 26637000 | 11000 |
| 6 | VITVarB40-14_v2.0.hap1.chr14 | CNV10605 | 26696001 | 26702000 | 6000 |
| 6 | VITVarB40-14_v2.0.hap1.chr14 | CNV10597 | 26524001 | 26528000 | 4000 |
| 6 | VITVarB40-14_v2.0.hap1.chr14 | CNV10602 | 26640001 | 26653000 | 13000 |
| 6 | VITVarB40-14_v2.0.hap1.chr14 | CNV10603 | 26655001 | 26686000 | 31000 |
| 6 | VITVarB40-14_v2.0.hap1.chr14 | CNV10604 | 26687001 | 26693000 | 6000 |
| 6 | VITVarB40-14_v2.0.hap1.chr14 | CNV10606 | 26701001 | 26716000 | 15000 |
| 6 | VITVarB40-14_v2.0.hap1.chr14 | CNV10607 | 26721001 | 26727000 | 6000 |
| 6 | VITVarB40-14_v2.0.hap1.chr14 | CNV10598 | 26530001 | 26611000 | 81000 |

| | | | | | |
|---|---|---|---|---|---|
| 6 | VITVarB40-14_v2.0.hap1.chr14 | CNV10608 | 26726001 | 26753000 | 27000 |
| 7 | VITVarB40-14_v2.0.hap1.chr15 | CNV10804 | 3295001 | 3300000 | 5000 |
| 7 | VITVarB40-14_v2.0.hap1.chr15 | CNV10808 | 3353001 | 3358000 | 5000 |
| 7 | VITVarB40-14_v2.0.hap1.chr15 | CNV10809 | 3359001 | 3364000 | 5000 |
| 7 | VITVarB40-14_v2.0.hap1.chr15 | CNV10810 | 3365001 | 3372000 | 7000 |
| 7 | VITVarB40-14_v2.0.hap1.chr15 | CNV10813 | 3431001 | 3438000 | 7000 |
| 7 | VITVarB40-14_v2.0.hap1.chr15 | CNV10814 | 3462001 | 3466000 | 4000 |
| 7 | VITVarB40-14_v2.0.hap1.chr15 | CNV10815 | 3467001 | 3471000 | 4000 |
| 7 | VITVarB40-14_v2.0.hap1.chr15 | CNV10800 | 3154001 | 3227000 | 20161 |
| 7 | VITVarB40-14_v2.0.hap1.chr15 | CNV10801 | 3227001 | 3233000 | 6000 |
| 7 | VITVarB40-14_v2.0.hap1.chr15 | CNV10802 | 3242001 | 3246000 | 4000 |
| 7 | VITVarB40-14_v2.0.hap1.chr15 | CNV10805 | 3304001 | 3316000 | 12000 |
| 7 | VITVarB40-14_v2.0.hap1.chr15 | CNV10806 | 3317001 | 3334000 | 17000 |
| 7 | VITVarB40-14_v2.0.hap1.chr15 | CNV10807 | 3340001 | 3344000 | 4000 |
| 7 | VITVarB40-14_v2.0.hap1.chr15 | CNV10812 | 3419001 | 3430000 | 11000 |
| 7 | VITVarB40-14_v2.0.hap1.chr15 | CNV10803 | 3262001 | 3282000 | 20000 |
| 7 | VITVarB40-14_v2.0.hap1.chr15 | CNV10811 | 3373001 | 3417000 | 44000 |
| 8 | VITVarB40-14_v2.0.hap1.chr15 | CNV11365 | 19204001 | 19209000 | 5000 |
| 8 | VITVarB40-14_v2.0.hap1.chr15 | CNV11364 | 19101001 | 19106000 | 5000 |

**Table S4.**
Name, sequence, p-values and adjusted p-values of the 115 significant kmers. P-values are based on associations with *X. fastidiosa* load (CFU/ml) using GEMMA. Adjusted p-values are Bonferroni corrected.

| kmer ID | kmer sequence | p-value | Adj. p-value |
|---|---|---|---|
| kmer1 | CATTTATTGCTTCTTAAAATGTAGAAACAAG | 6.03E-14 | 5.83E-05 |
| kmer2 | ATTTATTGCTTCTTAAAATGTAGAAACAAGG | 6.03E-14 | 5.83E-05 |
| kmer3 | TCCTTGTTTCTACATTTTAAGAAGCAATAAA | 6.03E-14 | 5.83E-05 |
| kmer6 | ACATTTATTGCTTCTTAAAATGTAGAAACAA | 1.92E-13 | 0.000185722 |
| kmer7 | TACATTTATTGCTTCTTAAAATGTAGAAACA | 1.92E-13 | 0.000185722 |
| kmer20 | TCTACATTTTAAGAAGCAATAAATGTATAAA | 8.10E-13 | 0.000783724 |
| kmer21 | ATACATTTATTGCTTCTTAAAATGTAGAAAC | 8.10E-13 | 0.000783724 |
| kmer22 | TTATACATTTATTGCTTCTTAAAATGTAGAA | 8.10E-13 | 0.000783724 |
| kmer23 | ACATTTTAAGAAGCAATAAATGTATAAAAGA | 8.10E-13 | 0.000783724 |
| kmer24 | CTACATTTTAAGAAGCAATAAATGTATAAAA | 8.10E-13 | 0.000783724 |
| kmer25 | CATTTTAAGAAGCAATAAATGTATAAAAGAA | 8.10E-13 | 0.000783724 |
| kmer26 | CTTTTATACATTTATTGCTTCTTAAAATGTA | 8.10E-13 | 0.000783724 |
| kmer27 | TATACATTTATTGCTTCTTAAAATGTAGAAA | 8.10E-13 | 0.000783724 |
| kmer39 | AAGTATTTCTAACTCTAATATTTTGGAATAA | 3.50E-12 | 0.003383923 |
| kmer40 | AGTATTTCTAACTCTAATATTTTGGAATAAT | 3.50E-12 | 0.003383923 |
| kmer43 | CACATCTCGAAGACTCGCCAAAAAATAAATA | 7.53E-12 | 0.007281675 |
| kmer44 | ACATCTCGAAGACTCGCCAAAAAATAAATAA | 7.53E-12 | 0.007281675 |
| kmer69 | ATTTTAAGAAGCAATAAATGTATAAAAGAAA | 3.62E-12 | 0.003504838 |
| kmer75 | CTCACATCTCGAAGACTCGCCAAAAAATAAA | 8.70E-12 | 0.008416304 |
| kmer85 | CAACCTTCTAAAATTAAGTTCCTGAAATAAA | 2.60E-11 | 0.025164858 |
| kmer86 | AACCTTCTAAAATTAAGTTCCTGAAATAAAT | 2.60E-11 | 0.025164858 |
| kmer95 | ATTTCTTTTATACATTTATTGCTTCTTAAAA | 9.96E-12 | 0.009632815 |
| kmer96 | AAGCAATAAATGTATAAAAGAAATTATACTT | 9.96E-12 | 0.009632815 |
| kmer97 | AAGAAGCAATAAATGTATAAAAGAAATTATA | 9.96E-12 | 0.009632815 |
| kmer98 | AGTATAATTTCTTTTATACATTTATTGCTTC | 9.96E-12 | 0.009632815 |
| kmer99 | AGCAATAAATGTATAAAAGAAATTATACTTA | 9.96E-12 | 0.009632815 |
| kmer100 | AATTTCTTTTATACATTTATTGCTTCTTAAA | 9.96E-12 | 0.009632815 |
| kmer101 | AGAAGCAATAAATGTATAAAAGAAATTATAC | 9.96E-12 | 0.009632815 |
| kmer102 | ATAATTTCTTTTATACATTTATTGCTTCTTA | 9.96E-12 | 0.009632815 |
| kmer103 | TAATTTCTTTTATACATTTATTGCTTCTTAA | 9.96E-12 | 0.009632815 |
| kmer104 | ATCTCGAAGACTCGCCAAAAAATAAATAAAT | 9.88E-12 | 0.009557721 |
| kmer105 | CATCTCGAAGACTCGCCAAAAAATAAATAAA | 9.88E-12 | 0.009557721 |
| kmer115 | ATTTATTTTTTGGCGAGTCTTCGAGATGTGA | 2.84E-11 | 0.027479841 |
| kmer126 | AAATATGAAGAAGAATATAAAAGACACAAGC | 1.99E-12 | 0.001924888 |

| | | | |
|---|---|---|---|
| kmer127 | AAAAATATGAAGAAGAATATAAAAGACACAA | 1.99E-12 | 0.001924888 |
| kmer128 | GGAAAAATATGAAGAAGAATATAAAAGACAC | 1.99E-12 | 0.001924888 |
| kmer129 | GAAAAATATGAAGAAGAATATAAAAGACACA | 1.99E-12 | 0.001924888 |
| kmer130 | AAAATATGAAGAAGAATATAAAAGACACAAG | 1.99E-12 | 0.001924888 |
| kmer131 | ACTCACATCTCGAAGACTCGCCAAAAAATAA | 4.26E-11 | 0.041168483 |
| kmer132 | TATACTCACATCTCGAAGACTCGCCAAAAAA | 4.26E-11 | 0.041168483 |
| kmer133 | TACTCACATCTCGAAGACTCGCCAAAAAATA | 4.26E-11 | 0.041168483 |
| kmer134 | TATATACTCACATCTCGAAGACTCGCCAAAA | 4.26E-11 | 0.041168483 |
| kmer135 | ATACTCACATCTCGAAGACTCGCCAAAAAAT | 4.26E-11 | 0.041168483 |
| kmer136 | ATATACTCACATCTCGAAGACTCGCCAAAAA | 4.26E-11 | 0.041168483 |
| kmer142 | TATTTAATTATTCCAAAATATTAGAGTTAGA | 3.12E-11 | 0.03016933 |
| kmer143 | ATTTAATTATTCCAAAATATTAGAGTTAGAA | 3.12E-11 | 0.03016933 |
| kmer144 | TTTAATTATTCCAAAATATTAGAGTTAGAAA | 3.12E-11 | 0.03016933 |
| kmer145 | TAATTATTCCAAAATATTAGAGTTAGAAATA | 3.12E-11 | 0.03016933 |
| kmer146 | ATTTCTAACTCTAATATTTTGGAATAATTAA | 3.12E-11 | 0.03016933 |
| kmer147 | AACTCTAATATTTTGGAATAATTAAATAATC | 3.12E-11 | 0.03016933 |
| kmer148 | CTAACTCTAATATTTTGGAATAATTAAATAA | 3.12E-11 | 0.03016933 |
| kmer149 | ATTATTTAATTATTCCAAAATATTAGAGTTA | 3.12E-11 | 0.03016933 |
| kmer156 | TCAACCTTCTAAAATTAAGTTCCTGAAATAA | 3.49E-11 | 0.033711675 |
| kmer159 | ACAGGAAAATGAGATGATTAGTGGGAAGGAG | 1.97E-11 | 0.019035426 |
| kmer160 | CTTCCCACTAATCATCTCATTTTCCTGTCAC | 1.97E-11 | 0.019035426 |
| kmer161 | CAGGAAAATGAGATGATTAGTGGGAAGGAGA | 1.97E-11 | 0.019035426 |
| kmer162 | GACAGGAAAATGAGATGATTAGTGGGAAGGA | 1.97E-11 | 0.019035426 |
| kmer163 | CCTTCCCACTAATCATCTCATTTTCCTGTCA | 1.97E-11 | 0.019035426 |
| kmer164 | ATATGTGCTTGAGCATTTTCATGTCGAAATC | 7.39E-12 | 0.007144371 |
| kmer165 | AATATGTGCTTGAGCATTTTCATGTCGAAAT | 7.39E-12 | 0.007144371 |
| kmer166 | CTTTGATTTCGACATGAAAATGCTCAAGCAC | 7.39E-12 | 0.007144371 |
| kmer167 | ACTTTGATTTCGACATGAAAATGCTCAAGCA | 7.39E-12 | 0.007144371 |
| kmer168 | GCTTGAGCATTTTCATGTCGAAATCAAAGTA | 7.39E-12 | 0.007144371 |
| kmer169 | TATGTGCTTGAGCATTTTCATGTCGAAATCA | 7.39E-12 | 0.007144371 |
| kmer170 | AATACTTTGATTTCGACATGAAAATGCTCAA | 7.39E-12 | 0.007144371 |
| kmer171 | ATACTTTGATTTCGACATGAAAATGCTCAAG | 7.39E-12 | 0.007144371 |
| kmer172 | TAATATGTGCTTGAGCATTTTCATGTCGAAA | 7.39E-12 | 0.007144371 |
| kmer173 | ATGTGCTTGAGCATTTTCATGTCGAAATCAA | 7.39E-12 | 0.007144371 |
| kmer174 | GAATACTTTGATTTCGACATGAAAATGCTCA | 7.39E-12 | 0.007144371 |
| kmer175 | TGTGCTTGAGCATTTTCATGTCGAAATCAAA | 7.39E-12 | 0.007144371 |
| kmer177 | GAGTGACAGGAAAATGAGATGATTAGTGGGA | 2.15E-11 | 0.020748614 |
| kmer178 | AGTGACAGGAAAATGAGATGATTAGTGGGAA | 2.15E-11 | 0.020748614 |
| kmer179 | ATGCTGGAGTGACAGGAAAATGAGATGATTA | 2.15E-11 | 0.020748614 |
| kmer180 | CACTAATCATCTCATTTTCCTGTCACTCCAG | 2.15E-11 | 0.020748614 |

| | | | |
|---|---|---|---|
| kmer181 | CTAATCATCTCATTTTCCTGTCACTCCAGCA | 2.15E-11 | 0.020748614 |
| kmer182 | CCCACTAATCATCTCATTTTCCTGTCACTCC | 2.15E-11 | 0.020748614 |
| kmer183 | CCACTAATCATCTCATTTTCCTGTCACTCCA | 2.15E-11 | 0.020748614 |
| kmer184 | ACTAATCATCTCATTTTCCTGTCACTCCAGC | 2.15E-11 | 0.020748614 |
| kmer198 | CAGGGAATACTTTGATTTCGACATGAAAATG | 1.61E-11 | 0.015604265 |
| kmer199 | AGGGAATACTTTGATTTCGACATGAAAATGC | 1.61E-11 | 0.015604265 |
| kmer200 | GAGCATTTTCATGTCGAAATCAAAGTATTCC | 1.61E-11 | 0.015604265 |
| kmer201 | CCTCAGGGAATACTTTGATTTCGACATGAAA | 1.61E-11 | 0.015604265 |
| kmer202 | CATGTCGAAATCAAAGTATTCCCTGAGGTCA | 1.61E-11 | 0.015604265 |
| kmer203 | CTCAGGGAATACTTTGATTTCGACATGAAAA | 1.61E-11 | 0.015604265 |
| kmer204 | AGCATTTTCATGTCGAAATCAAAGTATTCCC | 1.61E-11 | 0.015604265 |
| kmer205 | ATTTTCATGTCGAAATCAAAGTATTCCCTGA | 1.61E-11 | 0.015604265 |
| kmer206 | ATGTCGAAATCAAAGTATTCCCTGAGGTCAA | 1.61E-11 | 0.015604265 |
| kmer207 | ACCTCAGGGAATACTTTGATTTCGACATGAA | 1.61E-11 | 0.015604265 |
| kmer208 | GACCTCAGGGAATACTTTGATTTCGACATGA | 1.61E-11 | 0.015604265 |
| kmer264 | ACCTTGACCTCAGGGAATACTTTGATTTCGA | 3.05E-11 | 0.029516454 |
| kmer265 | CCTTGACCTCAGGGAATACTTTGATTTCGAC | 3.05E-11 | 0.029516454 |
| kmer266 | CGAAATCAAAGTATTCCCTGAGGTCAAGGTA | 3.05E-11 | 0.029516454 |
| kmer373 | GAATATAAAAGACACAAGCTGATTAGATAAA | 8.65E-12 | 0.008364752 |
| kmer374 | AATATAAAAGACACAAGCTGATTAGATAAAC | 8.65E-12 | 0.008364752 |
| kmer385 | AATATGAAGAAGAATATAAAAGACACAAGCT | 1.97E-11 | 0.019032593 |
| kmer425 | CAAAAAAATAGACAAATAAGAGGCAAAGCGG | 4.85E-11 | 0.046939693 |
| kmer491 | AGTTTATCTAATCAGCTTGTGTCTTTTATAT | 2.39E-11 | 0.023153186 |
| kmer512 | AAAGTAGGATACTCACATTTGATGACCAAAG | 1.96E-11 | 0.01892492 |
| kmer513 | CAAAAGTAGGATACTCACATTTGATGACCAA | 1.96E-11 | 0.01892492 |
| kmer514 | CCCAAAAGTAGGATACTCACATTTGATGACC | 1.96E-11 | 0.01892492 |
| kmer515 | GTCATCAAATGTGAGTATCCTACTTTTGGGA | 1.96E-11 | 0.01892492 |
| kmer516 | AAAAGTAGGATACTCACATTTGATGACCAAA | 1.96E-11 | 0.01892492 |
| kmer517 | CCAAAAGTAGGATACTCACATTTGATGACCA | 1.96E-11 | 0.01892492 |
| kmer518 | AAGTAGGATACTCACATTTGATGACCAAAGG | 1.96E-11 | 0.01892492 |
| kmer561 | AAGTTTATCTAATCAGCTTGTGTCTTTTATA | 2.81E-11 | 0.027166424 |
| kmer562 | CCAAGTTTATCTAATCAGCTTGTGTCTTTTA | 2.81E-11 | 0.027166424 |
| kmer563 | ATAAAAGACACAAGCTGATTAGATAAACTTG | 2.81E-11 | 0.027166424 |
| kmer564 | AAAGACACAAGCTGATTAGATAAACTTGGAA | 2.81E-11 | 0.027166424 |
| kmer565 | AAAAGACACAAGCTGATTAGATAAACTTGGA | 2.81E-11 | 0.027166424 |
| kmer644 | ACCTTCTTGGAATTTGAAAGAATATTGTCCA | 4.47E-11 | 0.043272587 |
| kmer645 | ATGGACAATATTCTTTCAAATTCCAAGAAGG | 4.47E-11 | 0.043272587 |
| kmer1218 | AGCTTGATGCAGTTGTCTTCTGAAAGCCATA | 2.32E-11 | 0.022438949 |
| kmer2408 | CTACAAATATCATCATCGATGGTATTCAATA | 4.16E-11 | 0.040252903 |
| kmer2409 | TACTACAAATATCATCATCGATGGTATTCAA | 4.16E-11 | 0.040252903 |

| | | | |
|---|---|---|---|
| kmer2410 | ACTACAAATATCATCATCGATGGTATTCAAT | 4.16E-11 | 0.040252903 |

**Table S5.**
Best hit mapping information of significant kmers to hap1, hap2 and unplaced contigs. See Excel file Table5.xlsx

**Table S6.**
Gene annotation of genes within PD-associated peaks. See Excel file TableS6.xlsx

**Table S7.**
Presence-absence matrix of significant kmers in *V. arizonica.* See Excel file TableS7.xlsx

**Table S8.**

Presence-absence matrix of significant kmers in other species. See Excel file TableS8.xlsx

**Table S9.**

Alignments and expression results for candidate genes from Aguero et al. (2022). See Excel file TableS9.xlsx
